# 2,3,7,8-Tetrachlorodibenzo-*p*-dioxin (TCDD) elicited dose-dependent shifts in the murine urinary metabolome associated with hepatic AHR-mediated differential gene expression

**DOI:** 10.1101/2024.10.22.619714

**Authors:** Warren J. Sink, Russell Fling, Ali Yilmaz, Rance Nault, Delanie Goniwiecha, Jack R. Harkema, Stewart F. Graham, Timothy Zacharewski

**Affiliations:** Michigan State University, Department of Biochemistry and Molecular Biology, East Lansing, MI 48823, USA; Michigan State University, Institute for Integrative Toxicology, East Lansing, MI 48824, USA; Corewell Health Research Institute, Royal Oak, MI 48073, USA; Oakland University-William Beaumont School of Medicine, Rochester, MI 48309, USA; Michigan State University, Department of Pharmacology and Toxicology, East Lansing, MI 48824, USA; Middlebury College, Neuroscience Faculty, 14 Old Chapel Rd, Middlebury, VT 05753, USA; Michigan State University, Pathobiology & Diagnostic Investigation, East Lansing, MI, United States of America

**Keywords:** 2,3,7,8-tetrachlorodibenzo-p-dioxin (TCDD), aryl hydrocarbon receptor (AHR), liver, toxicogenomics, 1-D ^1^H NMR, trimethylamine N-oxide (TMAO), glycolic acid, 1-methyl nicotinamide (1MN), histidine, branched-chain amino acid (BCAA)

## Abstract

Epidemiological evidence suggests an association between dioxin and dioxin-like compound (DLC) exposure and human liver disease. The prototypical DLC, 2,3,7,8-tetrachlorodibenzo-*p*-dioxin (TCDD), has been shown to induce the progression of reversible hepatic steatosis to steatohepatitis with periportal fibrosis and biliary hyperplasia in mice. Although the effects of TCDD toxicity are mediated by aryl hydrocarbon receptor (AHR) activation, the underlying mechanisms of TCDD-induced hepatotoxicity are unresolved. In the present study, male C57BL/6NCrl mice were gavaged every 4 days for 28 days with 0.03 - 30 μg/kg TCDD and evaluated for liver histopathology and gene expression as well as complementary 1-dimensional proton magnetic resonance (1D-^1^H NMR) urinary metabolic profiling. Urinary trimethylamine (TMA), trimethylamine *N*-oxide (TMAO), and 1-methylnicotinamide (1MN) levels were altered by TCDD at doses ≤ 3 μg/kg; other urinary metabolites, like glycolate, urocanate, and 3-hydroxyisovalerate, were only altered at doses that induced moderate to severe steatohepatitis. Bulk liver RNA-seq data suggested altered urinary metabolites correlated with hepatic differential gene expression corresponding to specific metabolic pathways. In addition to evaluating whether altered urinary metabolites were liver-dependent, published single-nuclear RNA-seq (snRNA-seq), AHR ChIP-seq, and AHR knockout gene expression datasets provide further support for hepatic cell-type and AHR-regulated dependency, respectively. Overall, TCDD-induced liver effects were preceded by and occurred with changes in urinary metabolite levels due to AHR-mediated changes in hepatic gene expression.

## INTRODUCTION

Metabolic dysfunction-associated steatotic liver disease (MASLD) includes a spectrum of liver pathologies, from simple and reversible hepatic steatosis to steatohepatitis with fibrosis in the absence of alcohol consumption, viral infection, or lipodystrophy that increases the risk for diabetes, end-stage liver disease, and hepatocellular carcinoma (HCC) (1–4). Genetics, lifestyle, and diet are commonly cited as causal factors. Yet, epidemiological studies have suggested MASLD and associated pathologies can also be linked to environmental pollutant exposures (5, 6). One such class of compounds, polychlorinated dibenzo-p-dioxins (PCDDs), such as the prototypical congener 2,3,7,8-tetrachlorodibenzo-*p*-dioxin (TCDD), and dioxin-like compounds (DLCs), are environmental pollutants, which have been associated with steatotic liver disease (SLD) and hepatotoxicity in humans (7–13).

TCDD toxicity is mediated by the ligand-activated transcription factor, the aryl hydrocarbon receptor (AHR) (14–16). The AHR is a member of the basic-helix-loop-helix (bHLH) Per-Arnt-Sim (PAS) transcription factor family (14, 15). Although it is promiscuous and binds many structurally diverse ligands with varying affinities, no single eminent physiological ligand has been identified (17, 18). Upon ligand binding, the AHR translocates to the nucleus where it heterodimerizes with the AHR nuclear translocator (ARNT). Canonically, the AHR-ARNT dimer binds to dioxin response elements (DREs) within the locus of target genes and recruits coactivators and RNA Polymerase II to elicit species-, sex-, age-, tissue-and cell-specific differential gene expression (14). However, AHR-mediated differential gene expression has also been shown to involve DRE-independent regulation (5, 19) and interactions with other transcription factors (20).

Although TCDD induces dose-dependent hepatic steatosis in mice that can progress to steatohepatitis with periportal fibrosis and biliary hyperplasia (13, 21, 22), the gene regulation of AHR-mediated hepatotoxicity from TCDD and DLC exposure is not wholly understood (23). Previous studies of TCDD have integrated complementary hepatic gene expression and metabolomics data to investigate the progression of SLD pathologies, including the roles of carbohydrate, lipid and amino acid metabolism disruption (21–32).

In this current study, we integrated complementary liver histopathology, hepatic transcriptomics, and 1-dimensional proton magnetic resonance (1D ^1^H NMR) analysis of urine to further investigate dose-dependent changes in metabolism following treatment with TCDD at levels that spanned background exposures to an intentional poisoning (22, 33). These data were supplemented by assessing the level and activity of select enzymes of interest, as well as the re-examination of published gene expression and metabolomics data. Urinary metabolites related to choline, glyoxylate, vitamin B3, and amino acid metabolism were dose-dependently altered by TCDD. Changes in trimethylamine, trimethylamine *N*-oxide (TMAO), and 1-methylnicotinamide (1MN) preceded moderate to severe liver steatohepatitis and were consistent with changes in hepatic gene expression. Other metabolites, such as glycolate, altered by TCDD were also associated with hepatic differential gene expression but observed only at doses that induced moderate to severe steatohepatitis. Collectively, altered urinary metabolite levels correlated with differential gene expression, suggesting changes were due to hepatic AHR activation by TCDD.

## RESULTS

### TCDD-Elicited Effects

The dose range and treatment regimen used in this study have been previously shown to elicit dose-and time-dependent liver pathologies in mice, including the progression of steatosis to steatohepatitis with periportal fibrosis and biliary hyperplasia (13, 21, 22, 34). Food consumption was not changed in any dose groups (Supplementary Figure 1) as previously reported (31). Euthanasia and tissue collection occurred between ZT 0-3 to control for changes due to diurnal regulation including oscillating liver weight (31, 35, 36). TCDD dose-dependently increased absolute and relative liver weights along with a decrease in body weight at 30 μg/kg (21, 31, 32). There was also a dose-dependent increase in hepatic steatosis, hepatocellular hypertrophy, mixed inflammatory cell infiltration (mainly monocytes and lymphocytes with lesser numbers of neutrophils), and biliary hyperplasia, especially at ≥10 μg/kg TCDD (Figure 1 and Table 1). A slight increase in hepatic necrosis (Table 1) was consistent with the modest increase in serum ALT (32). Overall, TCDD did not elicit overt systemic toxicity or hepatotoxicity but dose-dependently induced steatohepatitis with evidence of biliary hyperplasia at 30 μg/kg TCDD.

**Figure 1.**
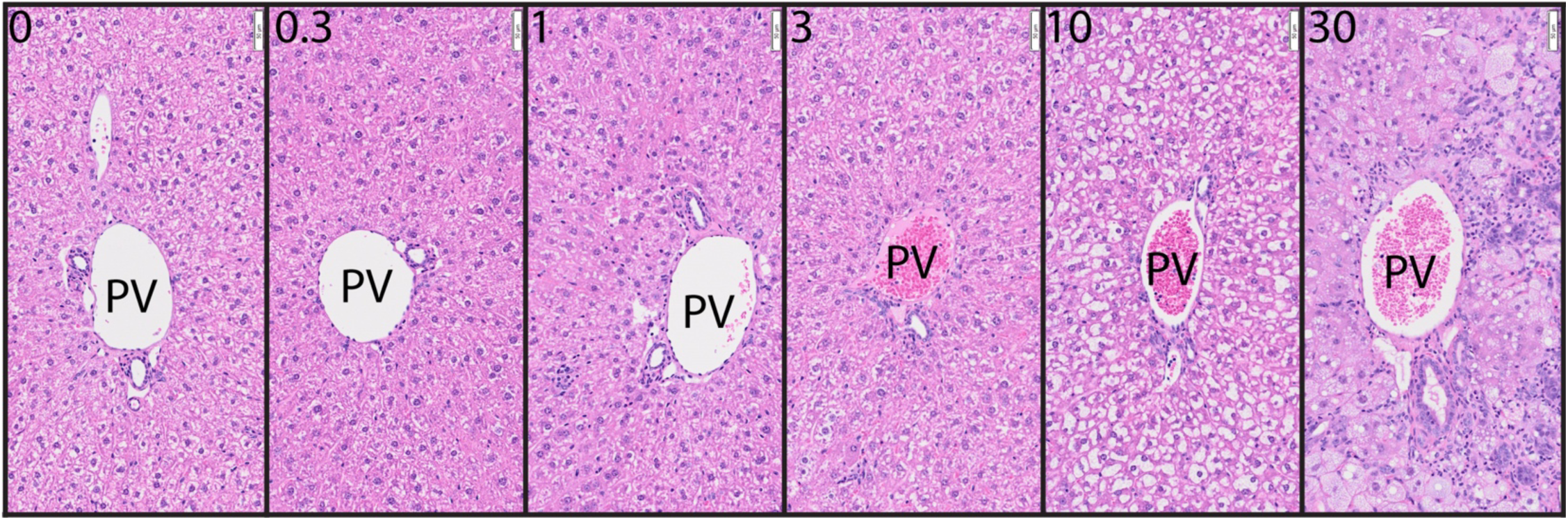
TCDD-elicited hepatic effects in C57BL/6NCrl malesgavaged every 4 days for 28 days. Hematoxylin and Eosin (H&E) staining of formalin-fixed liver sections from mice gavaged with 0 -30 μg/kg TCDD. The white scale bars (upper right hand corner of each image) indicate a length of 50 μm. Abbreviations: PV = portal vein.

**Table 1.**
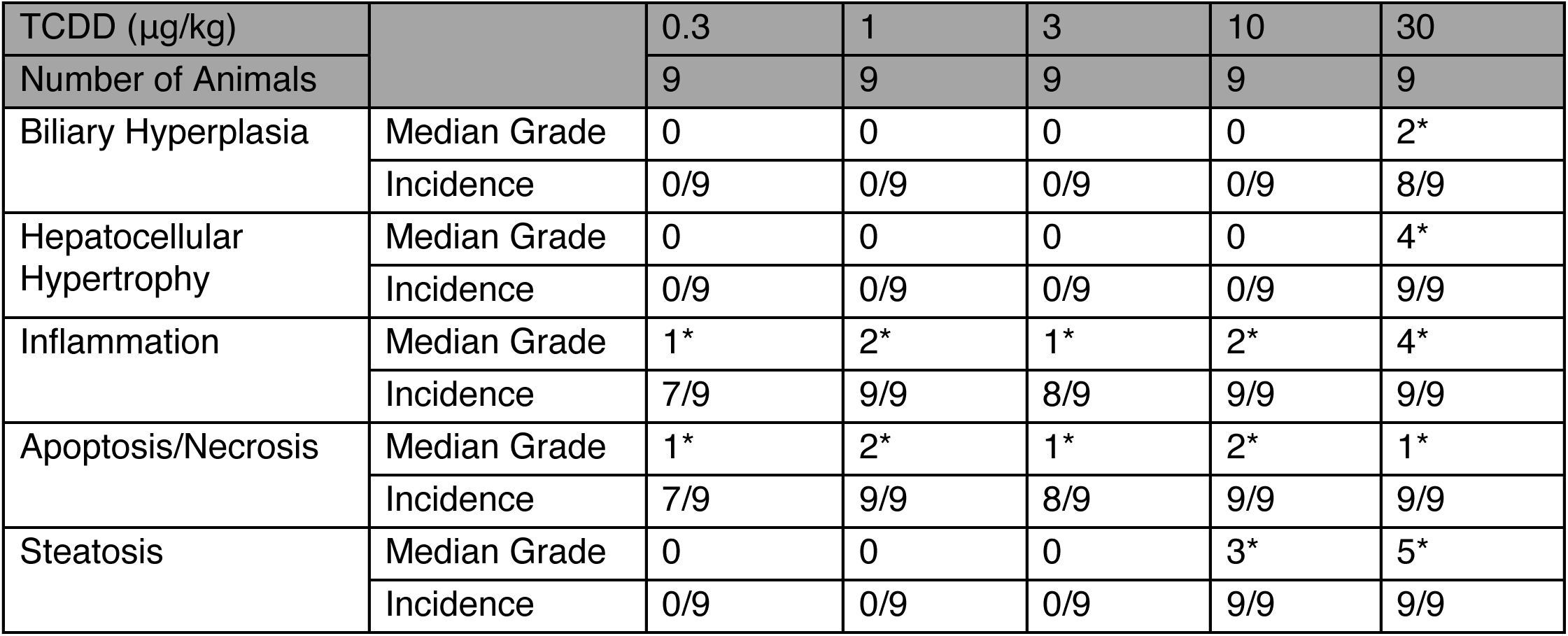
Semi-quantitative assessment of TCDD induced dose-dependent liver pathologies in male C57BL/6 mice. H&E-stained liver sections graded based on percent area using the following scale: 0 = 0%; 1 = minimal, less than 10%; 2 = mild, 10 to less than 25%; 3 = moderate, 25 to 50%; 4 = marked, 50 to less than 75%; 5 = severe, 75 to 100%. Median grade = median of histopathological grades. Incidence = occurrence of pathology in mice. An asterisk (*) indicates a significant (p ≤ 0.05) Dunnett’s test when compared to controls following a significant Kruskal-Wallis test.

| TCDD (µg/kg) |  | 0.3 | 1 | 3 | 10 | 30 |
| --- | --- | --- | --- | --- | --- | --- |
| Number of Animals |  | 9 | 9 | 9 | 9 | 9 |
| Biliary Hyperplasia | Median Grade | 0 | 0 | 0 | 0 | 2* |
|  | Incidence | 0/9 | 0/9 | 0/9 | 0/9 | 8/9 |
| Hepatocellular Hypertrophy | Median Grade | 0 | 0 | 0 | 0 | 4* |
|  | Incidence | 0/9 | 0/9 | 0/9 | 0/9 | 9/9 |
| Inflammation | Median Grade | 1* | 2* | 1* | 2* | 4* |
|  | Incidence | 7/9 | 9/9 | 8/9 | 9/9 | 9/9 |
| Apoptosis/Necrosis | Median Grade | 1* | 2* | 1* | 2* | 1* |
|  | Incidence | 7/9 | 9/9 | 8/9 | 9/9 | 9/9 |
| Steatosis | Median Grade | 0 | 0 | 0 | 3* | 5* |
|  | Incidence | 0/9 | 0/9 | 0/9 | 9/9 | 9/9 |

### 1-D ^1^H NMR Urinary Metabolite Profiling

In addition to TCDD-induced hepatic pathologies, treatment has been shown to cause AHR-mediated metabolic reprogramming and altered metabolite levels across various tissues. 1-D ^1^H NMR analysis of urine (*n* = 5 per dose) collected on post-natal day (PND) 55 (i.e., on day 25 of 28-day study; 1 day after the last gavage) between ZT 0 – 3 identified 107 unique metabolites in mouse urine samples (Supplementary Table 1).

Creatinine normalization (CN) is commonly used for evaluating the relative level of urinary metabolites as urinary creatinine is assumed to be constant within individuals over time and across individuals, but normalizing to creatinine alone may under-or overestimate the levels of metabolites of interest (37). Other methods for resolving the orders of magnitude difference in urinary metabolite levels, such as probabilistic quotient normalization (PQN) have also been developed for 1-D ^1^H NMR (38). Principal component analysis (PCA) was used to evaluate creatinine normalization, PQN, or the combination to identify the best approach to resolve the dose-dependent separation of urine metabolites along PC1. Non-normalized urinary creatinine did not show a dose-dependent trend (Supplementary Figure 2A). PC1 from non-normalized urinary metabolite profile data was dominated by noise that accounted for 47.5 percent explained variance (PEV), while PC2 showed dose-dependent separation of urinary profiles at ≥10 μg/kg TCDD (PEV = 8.2%) (Supplementary Figure 2B). Similarly, PCA analyses of urinary profiles after CN or PQN were dominated by outliers for PC1 (PEV_CN_ = 46.6 % and PEV_PQN_ = 24.4 %) while PC2 showed dose-dependent separation (Supplementary Figure 2C-D). CN of urinary profiles followed by PQN provided the best resolution along PC1 (PEV_CPQN_ = 15.48 %) with separation of samples occurring due to altered glycolate, pyruvate, succinate, urocanate, and hydroxyisovalerate (2HIV) at ≥ 10 μg/kg (Fig S2E). Therefore, statistical analyses of urinary metabolite levels were conducted on metabolite levels first normalized to creatinine and then subject to PQN.

The levels of succinate, methionine, trimethylamine (TMA), trimethylamine *N*-oxide (TMAO), and 1-methylnicotinamide (1MN) were altered below 10 μg/kg TCDD (Figure 2A). Other urinary metabolites were only altered at 10 or 30 μg/kg TCDD, including pyruvate, tyrosine, glutamic acid, 2-hydroxyisovalerate, 3-hydroxyisovalerate, 2-hydroxyvalerate, methylsuccinate, methylamine (MA), butyrate, isobutyrate, formate, and glycolate (Figure 2A). Of the altered metabolites, the median lower (BMDL_10%_) to upper bound (BMDU_10%_) benchmark dose responses ranged from 3.2 – 9.0 μg/kg. This included 1MN (BMDU_10%_: 2.3 μg/kg), *N,N’*-dimethylglycine (DMG) (BMDU_10%_: 6.8 μg/kg), isobutyrate (BMDU_10%_: 8.1 μg/kg), tyramine (BMDU_10%_: 4.70 μg/kg), butyrate (BMDU_10%_: 8.79 μg/kg), TMAO (BMDU_10%_: 8.72 μg/kg), glycolate (BMDU_10%_: 9.26 μg/kg), 2HIV (BMDU_10%_: 8.06 μg/kg), and urocanate (BMDU_10%_: 8.34 μg/kg) (Figure 2B).

**Figure 2.**
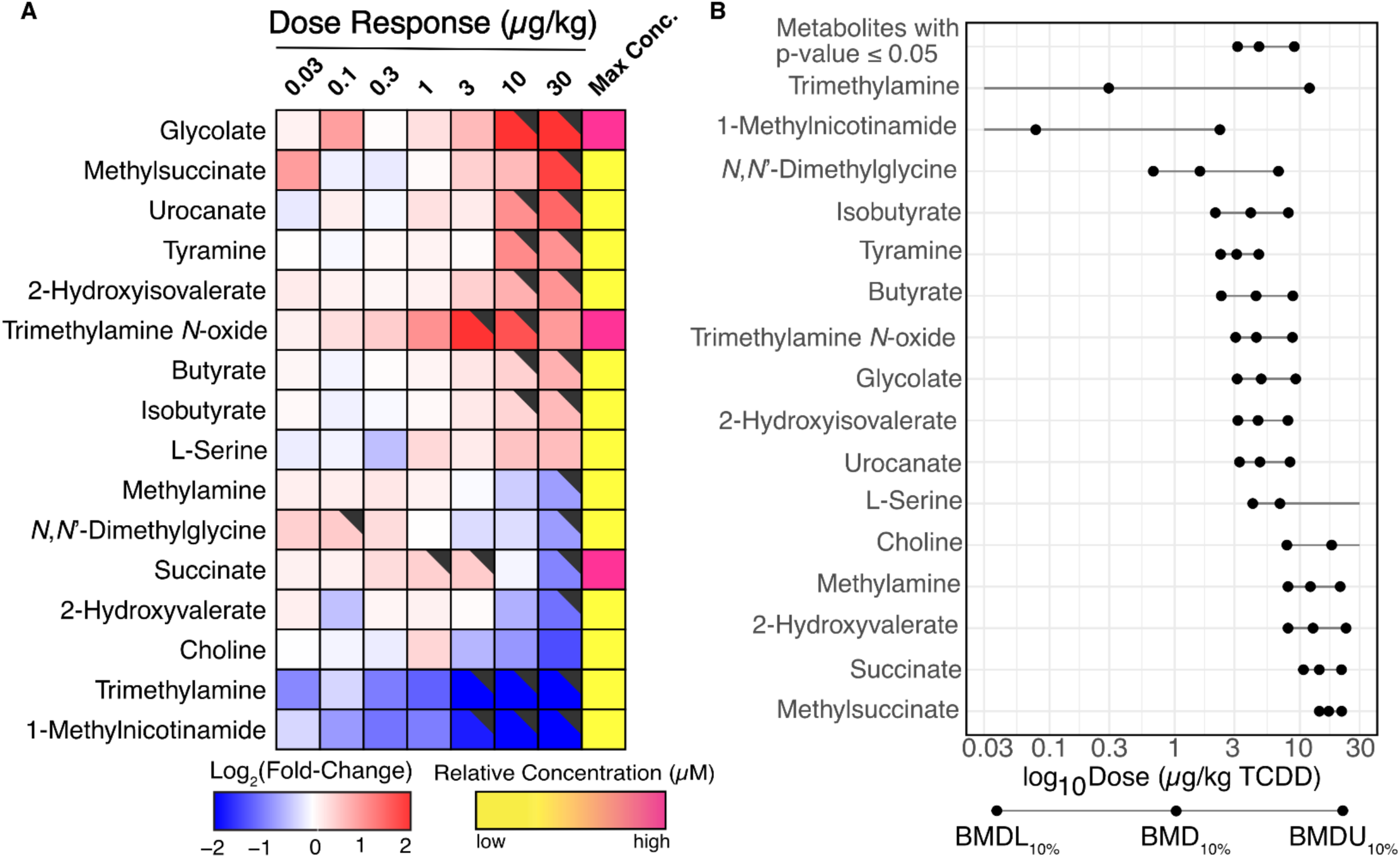
TCDD-elicited changes in urinary metabolites. Urine was collected from individual mice (n = 5) one day before euthanasia on PND 57 between ZT0-3. Metabolites were normalized by creatinine levels followed by probabilistic quotient normalization. (A) Heatmap of dose-dependent changes in metabolite levels. The black triangle (upper right hand corner) indicates a significant (p ≤ 0.05) change from controls following Benjamini-Hochberg (BH) and *post hoc* Dunnett’s tests relative to urine from sesame oil gavaged mice. Maximum concentrations are colored according to low (0.25 μM / μM creatinine) and high concentrations (2 μM / μM creatinine). (B) Benchmark dose-response values for metabolites in (A) calculated using BMDExpress2 (v2.3). The 10% benchmark dose (BMD_10%_) was estimated with a 95% confidence interval to calculate a lower (BMDL_10%_) and an upper (BMDU_10%_) bound. The median BMDL_10%_, BMD_10%_, and BMDU_10%_ of all significant metabolites (p-value ≤ 0.05).

### Integration of Hepatic Gene Expression

To identify metabolic pathways responsible for the altered urinary metabolite levels following TCDD treatment, liver samples (n = 5 per dose) from the same study were examined by bulk liver RNA-seq. TCDD caused a dose-dependent shift of the bulk hepatic transcriptome at ≥ 1 μg/kg TCDD (Figure 3A). Approximately, 6,722 genes in total were differentially expressed (DEGs) by at least one dose (Figure 3B) (Supplementary Table 2). 2,983 DEGs had a BMD_10%_ upper bound (BMDU_10%_) < 10 μg/kg TCDD (Figure 3C), suggesting widespread differential gene regulation below the dose at which TCDD elicited marked steatohepatitis, fibrosis, and biliary hyperplasia. Functional analysis of DEGs below 10 μg/kg TCDD showed enrichment for metabolic pathways associated with TCDD-altered urinary metabolite levels, including the metabolism of pyruvate (KEGG:00620), choline (KEGG:05231), branched-chain amino acids (BCAAs) (KEGG:00280), glyoxylate (KEGG:00630), and tryptophan (KEGG:00380) (Figure 3D).

**Figure 3.**
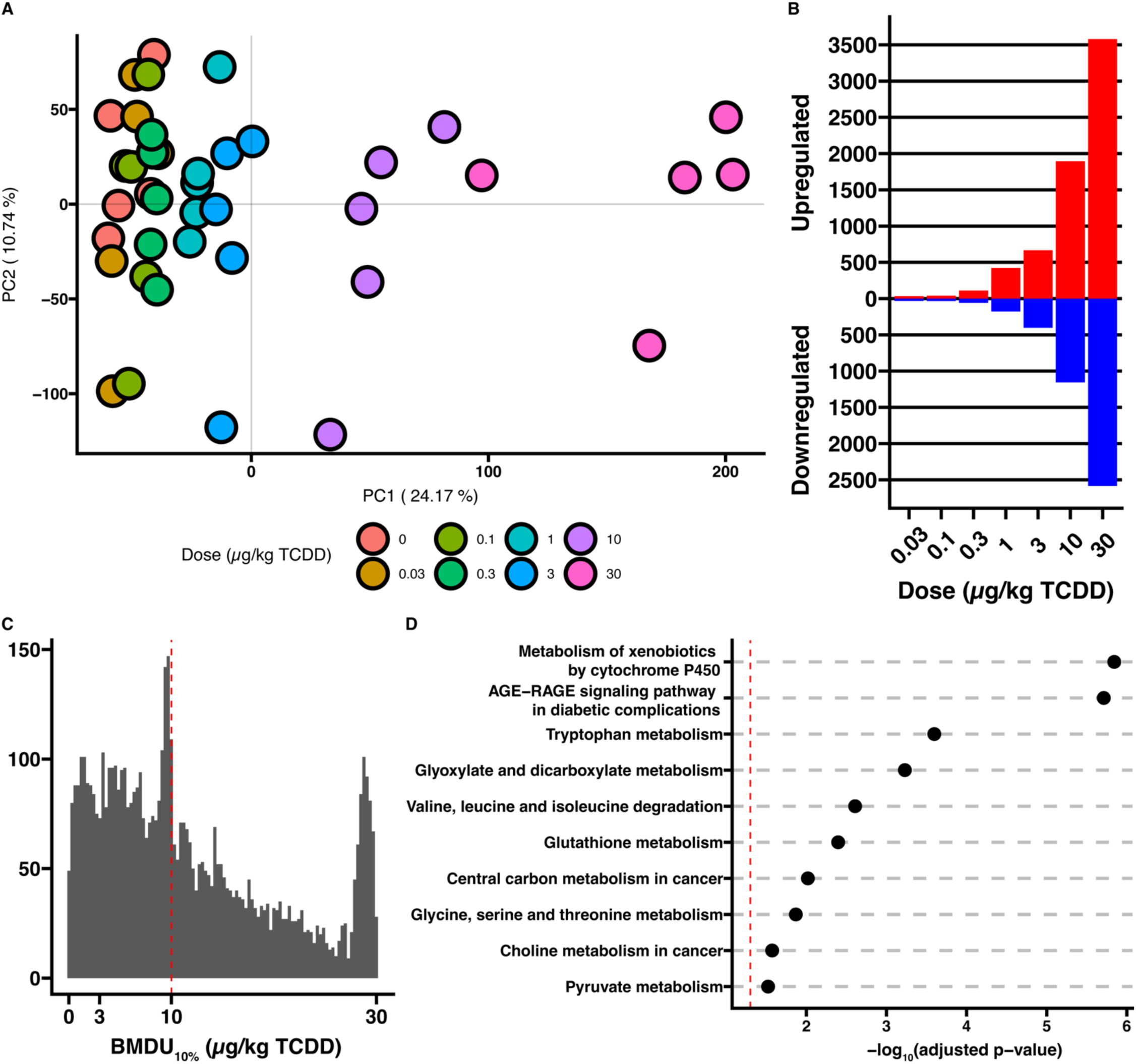
TCDD elicited dose-dependent bulk liver differential gene expression analysis. Mice gavaged every 4 days for 28 days with 0.03 – 30 μg/kg TCDD or sesame oil vehicle were euthanized on PND 58 between ZT0-3. Bulk liver RNA-seq was performed (n = 5). (A) Principal component analysis of all genes expressed in the liver. (B) Number of differentially expressed (IFold-ChangeI ≥ 1.5 and a P1(t) ≥ 0.8) induced and repressed genes at each dose. (C) The 10% benchmark dose (BMD_10%_) of all differentially expressed genes was estimated with a 95% confidence interval to calculate a lower (BMDL_10%_) and an upper (BMDU_10%_) bound using BMDExpress2 (v2.3). The red dashed line indicates 10 μg/kg TCDD. (D) KEGG pathways enriched for DEGs with a BMDU_10%_ less than 10 μg/kg TCDD. The dashed red line indicates -log_10_(adjusted p-value = 0.05).

### Choline Metabolism

Choline and carnitine are vitamin-like nutrients, which may be metabolized by gut microbiota to form TMA prior to their absorption (39) (Figure 4A and Supplementary Figure 3A). Urinary choline levels decreased 2.6-fold following TCDD treatment, while no change occurred in carnitine (Figure 4C). No significant change was detected in mouse cecum in the copy number of relevant TMA-producing microbial enzymes, including cutC (EC: 4.3.99.4), grdB (EC: 1.21.4.4), or cntA (EC: 1.14.13.139) (Supplementary Figure 3B-E). Moreover, as with previous reports (25, 26), there was little evidence suggesting choline was redirected to phosphatidylcholine (PtdC) biosynthesis by the Kennedy pathway or the synthesis of betaine to support homocysteine remethylation to methionine (Supplementary Figure 3A). Phosphorylcholine levels were unaltered (Supplementary Figure 3F), while *Chpt1*, which encodes the final and rate-limiting step of PtdC biosynthesis, was repressed 2.5-fold (Supplementary Figure 3G). *N*,*N*’-dimethylglycine (DMG) is produced when betaine donates a methyl group for the regeneration of methionine catalyzed by BHMT and was reduced 1.7-fold by TCDD (Supplementary Figure 3H). *Chdh* and *Aldh7a1*, which respectively encode enzymes for the two consecutive oxidations of choline è DMG è betaine (undetected in urine), were induced 1.3-fold and decreased 2.0-fold, respectively (Supplementary Figure 3I). Considering AHR-binding at pDREs and the 20-fold reduction in *Bhmt* (Supplementary Figure 3I), the decrease in urinary DMG was likely due to hepatic AHR-mediated downregulation.

**Figure 4.**
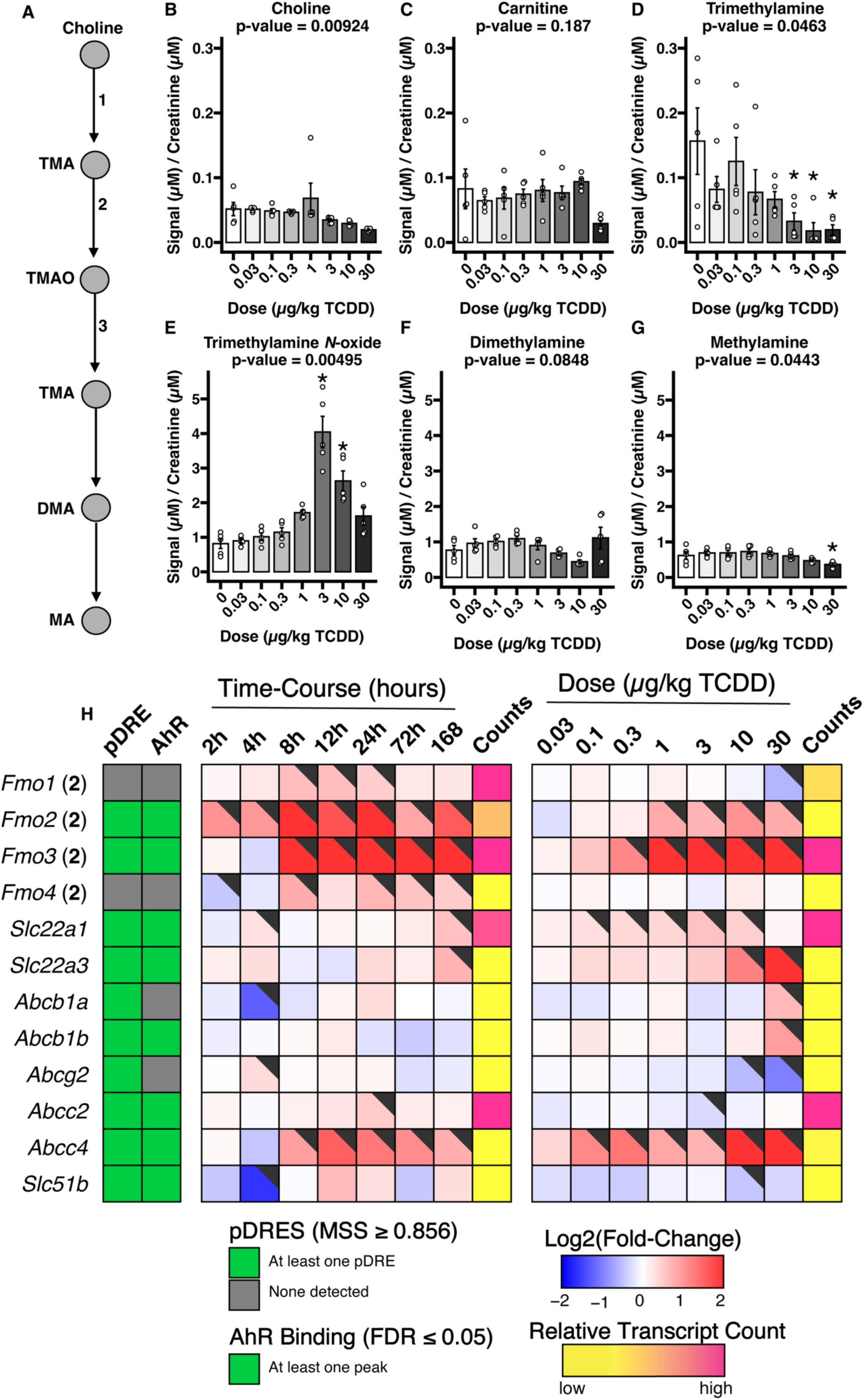
TCDD effect on urinary metabolites involved in choline metabolism. (A) The TMA pathway. Circles indicate metabolites. Arrows indicate enzymatic reaction. The numbers map to the genes in the heatmap (H), encoding the enzyme associated with the reaction. Abbreviations: TMA = trimethylamine; TMAO = trimethylamine *N*-oxide; DMA = dimethylamine; MA = methylamine. (B-G) Urinary metabolite levels as measured by 1-D ^1^HNMR (n = 5). An asterisk (*) indicates significance from a *post hoc* Dunnett test. (H) Heatmap of hepatic TMA/TMAO pathway related genes. pDREs were determined by a position weight matrix with a Matrix Similarity Score (MSS) ≥ 0.856. Hepatic AHR ChIP-seq was detected in mouse two hours after oral gavage with 30 μg/kg TCDD – green tiles indicate an FDR of ≤ 0.05 (GSE97634). In the dose response bulk RNA-seq gene expression, the black flags indicate a P1(t) ≥ 0.8. The Counts column refers to the maximum estimate of aligned reads of each gene where a lower level of expression (≤ 500 reads) is depicted in yelland a higher level of expression (≥ 10,000) is depicted in pink.

Metabolism of TMA in the gut and the liver leads to TMAO, dimethylamine (DMA), and methylamine with the possibility of TMA regeneration from TMAO (Figure 4A). Urinary TMA decreased 8.0-fold (Figure 4D), while TMAO increased as much as 5.0-fold at 3 μg/kg TCDD but fell to 2.0-fold at 30 μg/kg TCDD (Figure 4E). Urinary DMA was unchanged (Figure 4F), but methylamine was reduced 1.6-fold at 30 μg/kg TCDD (Figure 4G). FMO3 exhibits the greatest TMA oxidizing activity and is not normally expressed in adult male mice (40). However, 30 μg/kg TCDD induced *Fmo3* 5.3-fold at 8 hours, 102.6-fold after 7 days, and >600-fold after treatment every 4 days for 28 days (Figure 4H). Moreover, AHR genomic binding was present at a putative dioxin response element (pDRE) at the *Fmo3* transcription start site (TSS) 2 hours after a bolus gavage of TCDD (Figure 4H). *Fmo3* induction was greatest in hepatocytes as observed with single nuclear (sn)RNA-seq (Supplementary Figure 4A) with FMO3 protein levels also exhibiting dose-dependent increases (Supplementary Figure 4B). Urinary TMA and TMAO levels were altered at doses as low as 3 μg/kg TCDD coincident with the dose required to initiate induction of FMO3. Bulk liver RNA-seq datasets from whole body *Fmo3* knockout (KO) and wild-type (WT) mice treated with weekly injections of 25 μg/kg TCDD for 6 weeks concluded that *Fmo3* induction was responsible for the increased plasma TMAO levels although whether FMO3 induction is relevant to TCDD hepatotoxicity is questionable (Supplementary Figure 4C) (41, 42). Furthermore, whole-body *Ahr* KO mice treated with a bolus of TCDD confirmed *Fmo3* induction is AHR dependent (Supplementary Figure 4D-E) (43). Basal expression of *Fmo3* in adult male mice is minimal in all tissues except for lung. TCDD did not induce *Fmo3* in gonadal white adipose tissue, kidney, bone, or the intestinal tract (Supplementary Figure 4F). Therefore, increased FMO3 levels by TCDD in hepatocytes is likely the primary source of increased urinary TMAO levels.

Because TMAO is a charged compound, several potential transporters including organic cation transporters and ATP-binding cassettes have been reported to export or import TMAO across cell membranes (44). In TCDD-exposed mice, the most likely TMAO exporter from hepatocytes is ABCC4, which, located on the basolateral membrane, exports various molecules into the bloodstream. Its mRNA levels were induced 46.9-fold after oral gavage with 30 μg/kg TCDD every 4 days for 28 days, and the *Abcc4* locus exhibited AHR genomic binding in the presence of a pDRE. It exhibited similar gene expression to *Fmo3* in the time course with induction after a bolus TCDD dose as early as 8 hours post-gavage (Figure 4H).

### Hydroxyproline and glyoxal metabolism

Urinary glycolate, a hydroxyproline and glyoxal metabolism product (Figure 5A), was dose-dependently induced 7.4-fold by TCDD (Figure 5B), while the related compound, serine, was altered in the urine 1.6-fold (Figure 5C).

**Figure 5.**
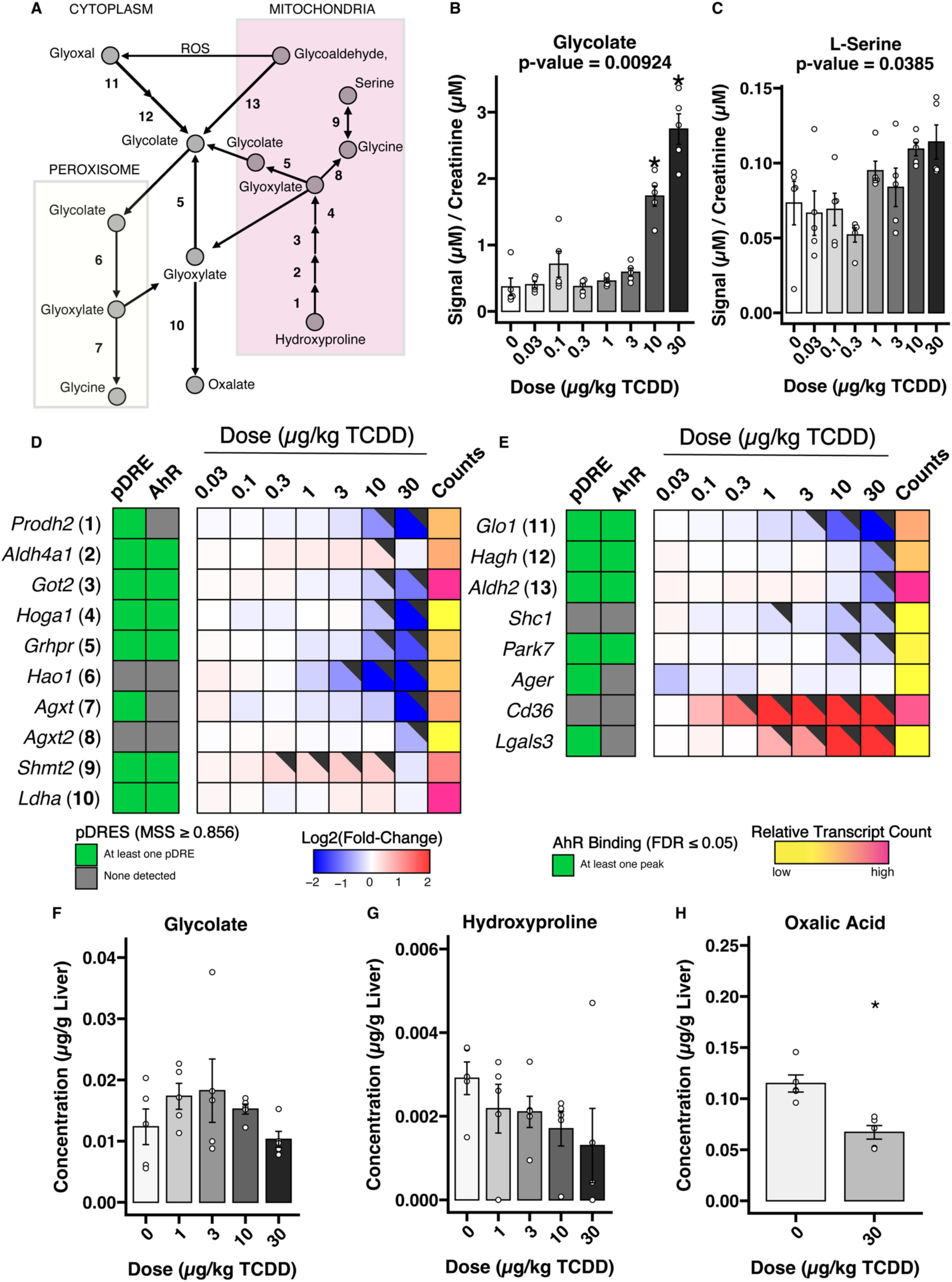
Hydroxyproline and glyoxal metabolism. (A) Hydroxyproline and glyoxal metabolism. (B – C) Urinary glycolate and L-serine levels measured by 1-D ^1^HNMR (n = 5). Data was normalized by creatinine and subsequently probabilistic quotient normalization. p-Values were calculated using a Kruskal-Wallis test and adjusted by the Benjamini-Hochberg method. An asterisk (*) indicates significance from a *post hoc* Dunnett test (p-value ≤ 0.05). (D – E) Heatmaps of hydroxyproline and glyoxal metabolism, respectively. pDREs were determined by a position weight matrix with a Matrix Similarity Score (MSS) cut off of ≥0.856 based on the sequence of characterized functional DREs. Hepatic AHR ChIP-seq was detected in mice two hours after oral gavage with 30 μg/kg TCDD (GSE97634). The green tiles indicate an FDR of ≤ 0.05. In the dose response bulk RNA-seq gene expression, the black flags indicate a P1(t) ≥ 0.8. The Counts column refers to the maximum estimate of aligned reads of each gene where a lower level of expression (≤ 500 reads) is depicted in yellow and a higher level of expression (≥ 10,000) is depicted in pink. (F – H) Hepatic metabolite levels measured by GC-MS (n = 5). p-Values were calculated using a Kruskal-Wallis test. An asterisk (*) indicates significance from a *post hoc* Dunnett test.

Hydroxyproline catabolism involves glyoxylate as an intermediate with subsequent metabolism to glycolate and oxalate (Figure 5A). In mice gavaged every 4 days for 28 days with TCDD, *Prodh2*, which encodes the first catalytic step of hydroxyproline metabolism, was downregulated 5.0-fold (Figure 5D). Subsequent reactions, encoded by *Got2* and *Hoga1*, which produce glyoxylate, were repressed and 4.0-fold, respectively (Figure 5D). *Grhpr*, which converts glyoxylate to glycolate, was also downregulated 2.7-fold (Figure 5D). In peroxisomes, HAO1 and AGXT convert glycolate into glycine. In the bulk liver RNA-seq data, *Hao1* and *Agxt* were downregulated 222.7-and 6.5-fold respectively (Figure 5D). Decreased expression of HAO1 and AGXT were confirmed by Western blot (Supplementary Figure 5A-B). Interestingly, no pDREs or AHR binding 2 hours after a bolus gavage of TCDD were detected at the *Hao1* locus (Figure 5D). snRNA-seq data suggests the dose-dependent repression of *Hao1* and *Agxt* primarily occurred in hepatocytes (Supplementary Figure 5C – D). Furthermore, *Hao1* expression was not affected in a TCDD-exposed whole-body *Ahr* KO model (Supplementary Figure E) (43).

Glycolate can also be formed by the detoxification of glyoxal (Figure 5A). Glyoxal is associated with advanced glycation end-products (AGEs) (45) unless it is metabolized into glycolate by *Glo1* and *Hagh*, which were downregulated 5.0-and 1.9-fold, respectively (Figure 5E). Reductions in GLO1 protein and activity were confirmed by western blot (Supplementary Figure 5F) and an enzymatic assay (Supplementary Figure 5G). *Glo1* and *Hagh* were primarily expressed in hepatocytes (Supplementary Figure 5H – I).

Although glycolate dose-dependently increased in the urine, hepatic glycolate and hydroxyproline levels were unchanged (Figure 5F – G), despite a slight reduction in hepatic oxalic acid (Figure 5H).

### Vitamin B3 metabolism

Urinary 1MN, a product of nicotinamide adenine dinucleotide (NAD^+^) and NAD phosphate (NADP^+^) metabolism (Figure 6A), is a biomarker of *de novo* NAD^+^ and NADP^+^ biosynthesis from tryptophan (46, 47). Although TCDD did not alter hepatic (30), serum (30), or urinary tryptophan levels (Figure 6B), 1MN was dose-dependently deceased 10.5-fold at 30 μg/kg TCDD (Figure 6C). Trigonelline also exhibited a decrease but did not achieve significance (Figure 6D). Indoleamine 2,3-dioxygenase 2 (*Ido2*) and tryptophan 2,3-dioxygenase (*Tdo2*) catalyze the first and rate-limiting step in the conversion of tryptophan to nicotinamide. Hepatic *Ido2* and *Tdo2* were repressed 4.0-and 1.6-fold, while *Ido1* expression was not detected (Figure 6E). The expression of genes encoding subsequent steps including *Afmid*, *Kmo*, *Kynu*, *Haao*, and *Qprt* were reduced 10.0-, 5.0-, 4.8-, 8.3-, and 6.3-fold, respectively (Figure 6E). *Ido2*, *Afmid*, *Kmo*, *Haao*, and *Qprt* all had AHR genomic binding in the presence of a pDRE (Figure 6E) suggesting repression may be AHR-mediated. *Nnmt*, which methylates nicotinamide to form 1-methylnicotinamide, was not repressed (Figure 6E).

**Figure 6.**
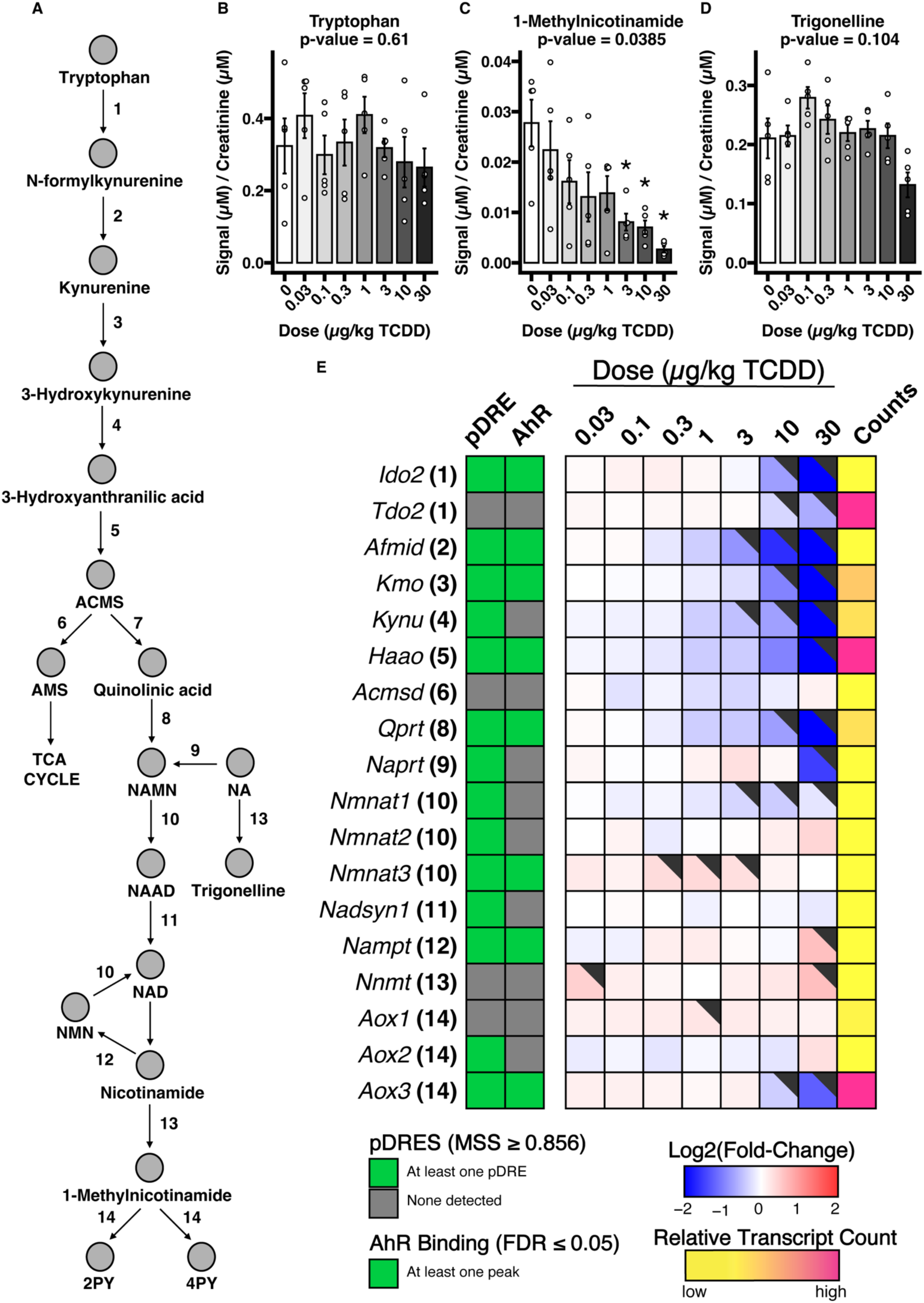
Effect of TCDD on nicotinamide biosynthesis. (A) Nicotinamide biosynthesis pathway. (B – D) Urinary tryptophan, trigonelline, and 1-methylnicotinamide levels as measured by 1-D ^1^H NMR (n = 5). Data was normalized by creatinine and subsequently probabilistic quotient normalization. An asterisk (*) indicates significance from a *post hoc* Dunnett test (p ≤ 0.05). (E) Heatmap of dose dependent effects of TCDD on gene expression associated with nicotinamide biosynthesis. pDREs were determined by a position weight matrix with a Matrix Similarity Score (MSS) cut off of ≥0.856 based on the sequence of characterized functional DREs. ChIP-seq analysis detected AHR genomic binding in mouse liver two hours after oral gavage with 30 μg/kg TCDD (GSE97634). The green tiles indicate an FDR of ≤ 0.05. In the dose response bulk RNA-seq gene expression, the black flags indicate a P1(t) ≥ 0.8. The Counts column refers to the maximum estimate of aligned reads of each gene where a lower level of expression (≤ 500 reads) is depicted in yellow and a higher level of expression (≥ 10,000) is depicted in pink. Abbreviations: ACMS = aminocarboxymuconate semialdehyde; AMS = aminomuconate semialdehyde; NAMN = nicotinic acid mononucleotide; NA = nicotinic acid; NAAD = nicotinic acid dinucleotide; NAD = nicotinamide dinucleotide; NMN = nicotinamide mononucleotide; 2PY = *N*-methyl-2-pyridone-5-carboxamide; 4PY = *N*-methyl-4-pyridone-3-carboxamide.

### Histidine metabolism

Histidine is metabolized by three possible pathways in mammalian liver, leading to the formation of histamine, imidazole pyruvate, or glutamate (Figure 7A). Urinary urocanate, an intermediate in histidine catabolism, was induced 2.8-fold (Figure 7B). Although urinary histidine was unchanged (Figure 7C) glutamate increased 1.7-fold (Figure 7D). TCDD has been shown to increase the levels of hepatic and serum histidine as well as hepatic glutamate (30). The first step in histidine metabolism is a nonoxidative deamination catalyzed by *Hal* (repressed 25.0-fold), which forms urocanate and ammonia (Figure 7E). In contrast, histidine decarboxylation by *Hdc* or conversion to imidazole propionate by *Hat1* were induced 1.6-and 1.8-fold, respectively (Figure 7E). Following *Hal*, *Uroc1*, which converts urocanate to imidazole propionate, was downregulated 10.0-fold (Figure 7E). Then, *Amdhd1*, which converts imidazole propionate to formiminoglutamate, and *Ftcd*, which converts formiminoglutamate to glutamate with the cofactor tetrahydrofolate (THF), were downregulated 7.7-and 5.3-fold, respectively (Figure 7E). Alternatively, histidine bioavailability may be modulated by the gut flora hut system (48). The hut genes detected by metagenomic analysis, hutH, hutU, and hutI, were not altered (Figure 7F-H), suggesting host metabolism was primarily responsible for altered histidine and urocanate levels.

**Figure 7.**
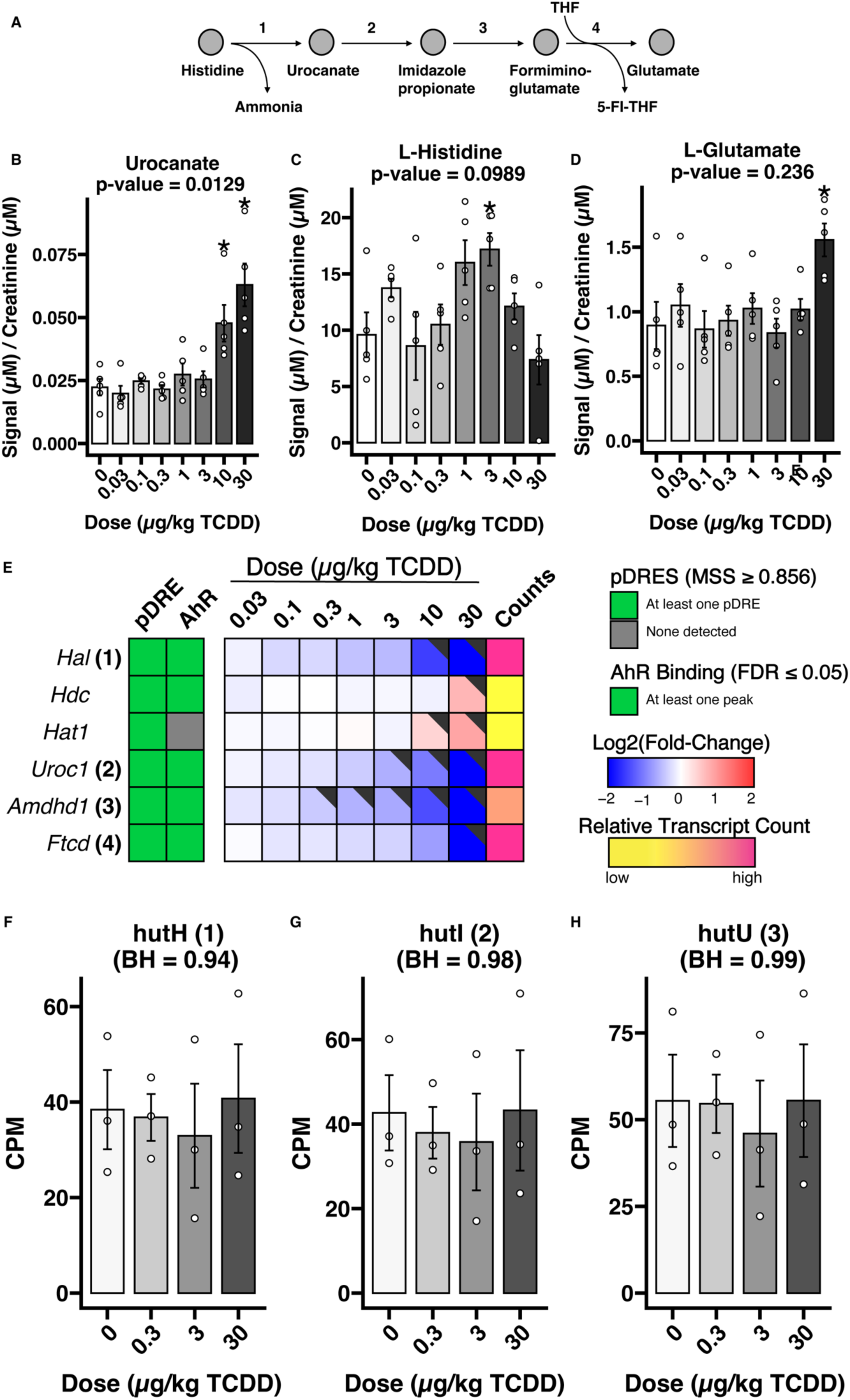
Histidine catabolism downregulation. (A) Histidine catabolism pathway. (B-D) 1-Dimensional ^1^H NMR signals of urinary histidine, urocanate, and glutamate normalized by creatinine and subsequently probabilistic quotient normalization (n = 5). An asterisk (*) indicates significance from a *post hoc* Dunnett test (p ≤ 0.05). (E) Heatmap of hepatic histidine metabolism. pDREs were determined by a position weight matrix with a Matrix Similarity Score (MSS) cut off of ≥0.856 based on the sequence of characterized functional DREs. ChIP-seq analysis detected AHR genomic binding in mouse liver two hours after oral gavage with 30 μg/kg TCDD (GSE97634). The green tiles indicate an FDR of ≤ 0.05. In the dose response bulk RNA-seq gene expression, the black flags indicate a P1(t) of at least 0.8. The Counts column refers to the maximum raw number of aligned reads to each gene where a lower level of expression (≤500 reads) is depicted in yellow and a higher level of expression (≥10,000) is depicted in pink. (F-H) Metagenomic copies per million (CPM) for all detected histidine utilization hub genes.

### Branched-chain amino acid catabolism

At homeostasis, catabolism of the branched-chain amino acids (BCAA) isoleucine, leucine, and valine share the first two enzymatic steps (Figure 8A and Supplementary Figure 7). Cytosolic BCAT1 or mitochondrial BCAT2 reversibly convert BCAA and α-ketoglutarate into branched-chain α-keto acids (BCKA) and glutamate (Figure 8A). Next, an irreversible oxidative decarboxylation by the branched-chain alpha-keto acid dehydrogenase (BCKDH) converts the BCKA into a coenzyme A (CoA)-activated branched hydrocarbon (Figure 8A). Although liver, serum, and urinary valine, leucine, and isoleucine were unaltered by TCDD (Figure 8B-D) (49), dose-dependent increases in the urinary levels of 2HIV, 3-hydroxyisovalerate (3HIV), and methylsuccinate (MSA) of 2.1-, 3.6-, and 3.6-fold, respectively, suggested BCAA metabolism was disrupted (Figure 8E-G), specifically within the BCKDH complex. In rodents and humans, the initial step of BCAA metabolism is extrahepatic (50), and therefore the modest effects of TCDD on hepatic *Bcat1* and *Bcat2* expression are negligible (Figure 8H). Moreover, TCDD had minor effects on *Bcat1* and *Bcat2* expression in muscle, kidney, gonadal white adipose tissue, and intestinal data (Supplementary Figure 6A – B). The second step of BCAA metabolism is catalyzed by BCKDH, an inner mitochondrial membrane multienzyme complex consisting of three enzymatic catalytic components: E_1_, encoded by *Bckdha* and *Bckdhb*; E_2_, encoded by *Dbt*; and E_3_, encoded by *Dld*. *Bckdha* was repressed 2.5-fold at 30 μg/kg TCDD in the absence of AHR genomic binding (Figure 8H). Furthermore, *Bckdhb* exhibited dose-dependent repression and AHR genomic binding in the presence of a pDRE (Figure 8H). E_2_ and E_3_, encoded by *Dbt* and *Dld*, respectively, were unaltered by treatment (Figure 8H). *Bckdha* and *Bckdhb* repression occurred mainly in the liver (Supplementary Figure 6C – D). BCKDH is inhibited by phosphorylation from the kinase BCKDK and is activated by the phosphatase PPM1K (Figure 8A). *Ppm1k* expression was dose-dependently reduced 2.9-fold, while *Bckdk* expression was unaltered. If BCKDH activity is inhibited, alternative metabolism of the BCKA occurs via BCKA-specific enzymes (Figure 8A). Like oxalic acid, 2HIV is produced from *Ldha* (51), the expression of which was unaltered by TCDD (Figure 8H). *Hpd* expression, which converts α-ketoisocaproate into 3HIV (52), was reduced 2.6-fold (Figure 8H). Although methylsuccinate is likely produced by the transformation of α-ketoisovalerate, the responsible enzyme is not known (53). *Bcat2*, *Bckdha*, *Bckdhb*, *Ppm1k*, and *Bckdk* were mainly expressed in hepatocytes and immune cells (Supplementary Figure 6E – I). KLF15, was repressed 2.3-fold (Supplementary Table 2), is a transcriptional regulator of BCAA metabolism (54). At homeostasis, KLF15 binds to the *Bckdha, Bckdhb, and Ppm1k* transcription start sites (TSS) (Supplementary Figure 7B – D). AHR genomic binding occurs at the TSS in the presence of a pDRE following TCDD treatment (Supplementary Figure 7A – C). Consistent with the repression of BCKDH, dose-dependent decreases in the hepatic levels of α-isobutyryl-CoA were identified in a complementary untargeted metabolomic dataset (32). In the same dataset, isovaleryl-CoA and α-methylbutyryl-CoA (also known as 2-methylbutyryl CoA) also exhibited dose-dependent decreases, but they could not be resolved from isomers (Supplementary Table 3). This untargeted mass spectrometry data was not verified by targeted analysis.

**Figure 8.**
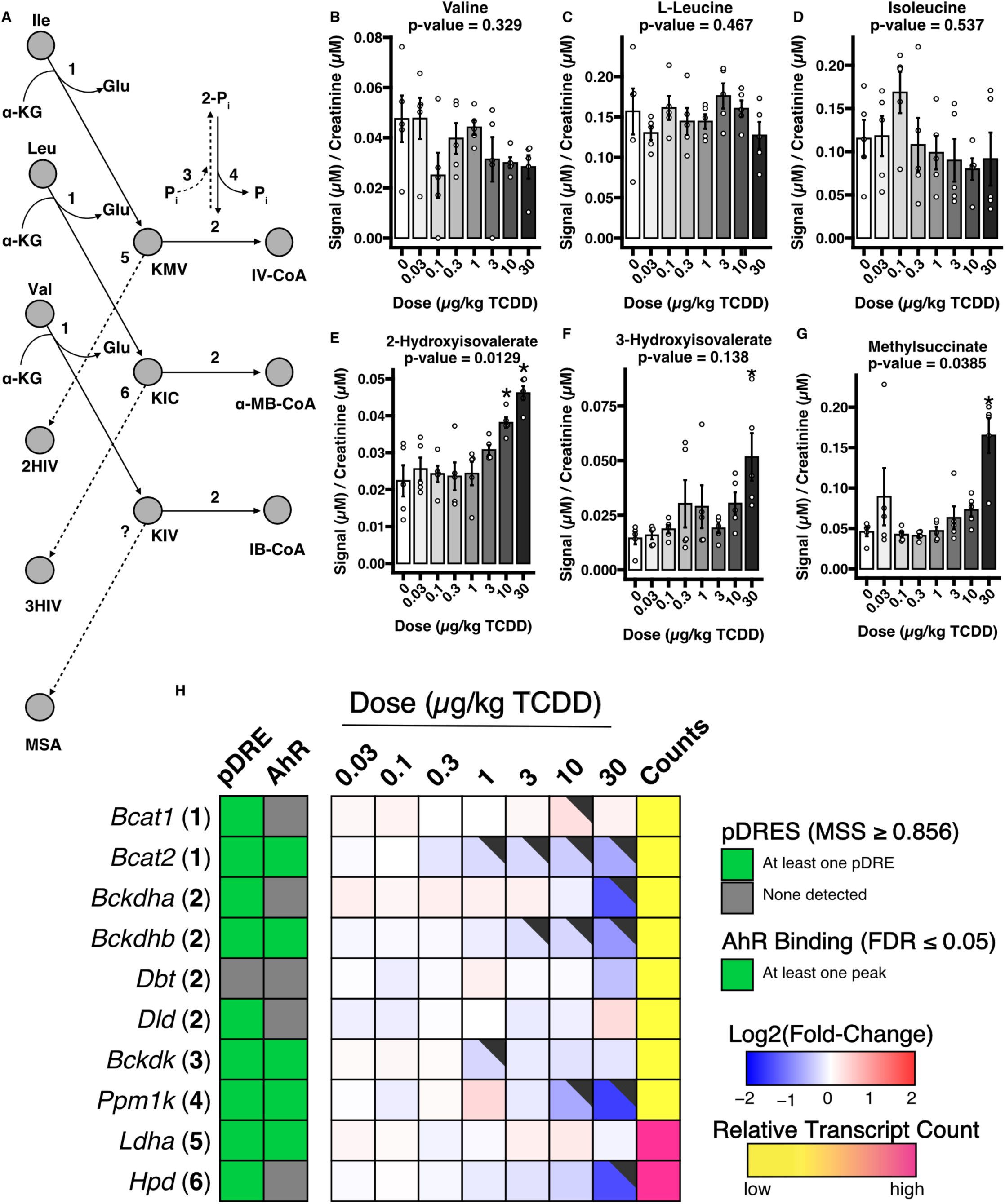
Alternative Branched-Chain Amino Acid Catabolism. (A) The first two steps in the canonical branched-chain amino acid (BCAA) catabolic pathway (bold lines). An alternative pathway (dashed lines) is also depicted when step 2 is inhibited. Step 2 may be inhibited by phosphorylation and activated by dephosphorylation (steps 3 and 4, respectively). The numbers map to the genes in the heatmap (H), encoding the enzyme associated with the reaction. (B-G) Urinary metabolite levels as measured by 1-D ^1^H NMR (n = 5). An asterisk (*) indicates significance from a post hoc Dunnett test. (H) Heatmap of hepatic BCAA catabolism related genes. pDREs were determined by a position weight matrix with a Matrix Similarity Score (MSS) ≥ 0.856. AHR ChIP-seq was detected in mouse liver two hours after oral gavage with 30 μg/kg TCDD – green tiles indicate an FDR of ≤ 0.05 (GSE97634). In the dose response bulk RNA-seq gene expression, the black flags indicate a P1(t) ≥ 0.8. The Counts column refers to the maximum estimate of aligned reads of each gene where a lower level of expression (≤ 500 reads) is depicted in yellow and a higher level of expression (≥ 10,000) is depicted in pink. Abbreviations: Ile = isoleucine; Leu = leucine; Valine = valine; αKG = α-ketoglutarate; Glu = glutamate; KMV = α-keto-β-methylvalerate; KIC = α-ketoisocaproate; KIV = α-ketoisovalerate; P_i_ = phosphate; IV-CoA = isovaleryl-CoA; α-MB-CoA = alpha-methyl-butyryl-CoA; IB-CoA = isobutyryl-CoA; 2HIV = 2-hydroxyisovalerate; 3HIV = 3-hydroxyisovalerate; MSA = methylsuccinate.

## DISCUSSION

Although TCDD and DLCs are associated with human and rodent liver dysfunction, the mechanism of TCDD hepatotoxicity via AHR-mediated metabolic reprogramming is not well understood (23). A recently published metabolome-wide association study analyzing the relation between exposure to DLCs in Dutch laborers identified changes in blood concentrations of certain metabolites, including those related to pyruvate, BCAA, histidine, methionine, and vitamin B3 metabolism (55). Similarly, the present study demonstrated that TCDD induced broad changes in murine urinary metabolite composition without altering daily food consumption, and these changes correlated with liver histopathology and differential gene expression. Urinary metabolites, such as 1MN, TMA, and TMAO, were altered at doses below those required to induce moderate to severe steatohepatitis (≤ 3 μg/kg), while pyruvate, glycolate, branched-chain amino acid catabolites, or urocanate, were only observed to be altered at ≥10 μg/kg TCDD.

Corroborating previous research (41, 56–59), AHR activation by TCDD induced FMO3 and increased urinary TMAO levels. The present study further demonstrated an inverse association with TMA and no alteration in the abundance of the primary microbial enzymes responsible for TMA production, including cutC. Individually, alteration of urinary TMA and TMAO levels occurred at doses as low as 3 μg/kg TCDD with *Fmo3* induction occurring at even lower doses. Considering the ionic charge of TMAO, the nominal bile acid and glutathione transporter ABCC4 is likely responsible for exporting TMAO from hepatocytes into the systemic circulation, although other transporters cannot be excluded. Moreover, although TCDD slightly reduced urinary choline, the mechanism by which this occurred is unclear from the present data. In humans, increased TMAO or TMA levels are proposed to increase the risk of cardiovascular disease (60). In rodents, there is a positive correlation between FMO3-catalyzed TMAO generation and atherosclerosis, with accelerated lesion development by DLCs (56). Furthermore, increased circulating TMAO has been associated with steatosis and all-cause mortality in people with steatosis (61, 62). This suggests TMAO may be a sensitive indicator of TCDD-induced steatotic development.

Glycolate is a relatively understudied two-carbon metabolite, normally found in mammals with little known about its role in homeostasis beyond being a precursor in hydroxyproline catabolism, glyoxal detoxification, and glyoxylate detoxification. Hydroxyproline is a significant component of collagen fibers and is elevated in fibrotic liver samples (63–66). Although bulk liver RNA-seq and AHR ChIP-seq data suggested AHR-mediated downregulation of the hydroxyproline catabolism, GC-MS analysis of aqueous liver extracts showed no convincing trend in hydroxyproline levels. A previous study using the same dosing regimen and study design reported mild periportal fibrosis, which may not be sufficient to detect increased hydroxyproline levels despite the induction of several collagen genes (22, 31). Dysregulation of glyoxylate metabolism can lead to kidney stones due to calcium oxalate accumulation (67, 68). Glyoxylate is reduced to glycolate by cytosolic or mitochondrial GRHPR. The glycolate may then be further metabolized by the peroxisomal HAO1 and AGXT to prevent calcium oxalate accumulation and the formation of kidney stones. In this study, *Hao1* and *Agxt* were repressed with corresponding reductions in protein levels. *Hao1* knockouts in rodents and dysfunction in humans are reported to increase glycolate levels compared to wild-type animals (69), implicating *Hao1* repression in elevated urinary glycolate levels.

Another possible source of glycolate, is glyoxal, an electrophilic dialdehyde that covalently binds to macromolecules, including lipids, proteins, and DNA, in a process referred to as glycation to form advanced glycation end-products (AGEs), a biomarker for diabetes (70). Glyoxal can be detoxified into glycolate by the glyoxalase system, specifically, GLO1 and HAGH using glutathione as a cofactor. *Glo1* and *Hagh* were both dose-dependently repressed by TCDD, with GLO1 activity also reduced. Overall, elevated urinary glycolate levels were most likely due to AHR-mediated repression of hepatic HAO1 expression. HAO1 repression may reflect peroxisomal metabolic reprogramming, including the loss of peroxisomal fatty acid metabolism, and even lower peroxisome levels, which may contribute to hepatic lipid accumulation.

Niacin, or vitamin B3, is a vitamin-like nutrient that can be replenished by the transformation of tryptophan. Diets deficient in tryptophan and niacin can be rescued with tryptophan administration (71). 1MN is a catabolite of niacin metabolism and correlates with *de novo* niacin production from tryptophan (46, 47). TCDD did not affect the liver, serum, or urinary tryptophan, but 1MN was dose-dependently reduced, suggesting a disruption in the *de novo* synthesis pathway. Formation of 1MN requires *S*-adenosylmethionine (SAM), which is dose-dependently reduced by TCDD (26). However, TCDD-induced decreases in 1MN may also be due to AHR-mediated repression of *Afmid* and other genes in the kynurenine pathway (72). Although decreased NAD^+^ is reported in liver disease (Guarino Metabolites 2019), gavaging mice every 4 days for 28 days with 30 μg/kg TCDD led to reduced NADP^+^ but did not lead to reduced levels of NAD^+^. Overall, further studies are needed to determine the role of *de novo* niacin production and SAM on lower 1MN levels and how this may affect kynurenine and NAD^+^.

Valine, leucine, and isoleucine are BCAAs that constitute a disproportionately large percentage of proteins (73). The first step of BCAA metabolism is catalyzed by branched-chain aminotransferases (BCAT) that occur in all tissues (50). The subsequent irreversible step by branched-chain keto acid dehydrogenase (BCKDH) primarily occurs in the liver and commits BCCAs to degradation for energy production (50). Urinary BCAA levels were altered by TCDD nor were the branched-chain keto-acids (BCKA) or coenzyme A-activated downstream products. Yet, the urinary levels of catabolic intermediates, 2-hydroxyisovalerate, 3-hydroxyisovalerate, and methylsuccinate, were increased by TCDD. All three are formed from different enzymes acting on respective BCKAs formed by BCAT metabolism. Moreover, a published metabolomics dataset for acyl-coenzyme A intermediates in liver extracts from mice gavaged every 4 days for 28 days with TCDD showed reciprocal decreases α-isobutyryl-CoA as well as the isomer unresolvable isovaleryl-CoA and/or α-methylbutyryl-CoA, suggesting inhibition of BCKDH activity. *Ppm1k*, a gene encoding a phosphatase, which prevents kinase-induced inhibition encoded by *Bckdk*, was also repressed. *Bckdk* and *Ppm1k* are regulated by KLF15, which was also repressed by TCDD. The repression of *Bckdha* and *Ppm1k* may lead to a decrease in total BCKDH and reduce the un-phosphorylated to phosphorylated BCKDH ratio, resulting in further inhibition of BCKDH, and increased formation of 2-hydroxyisovalerate, 3-hydroxyisovalerate, and methylsuccinate.

Although oxidative stress by Phase I enzyme induction contributes to AHR-mediated liver toxicity (74), accumulating evidence suggests TCDD-induced hepatotoxicity is due to the cumulative burden of the disruption of multiple metabolic pathways (23). Furthermore, the alteration of AHR-targeted metabolic pathways, which precede the development of liver pathology (*e.g.*, the TCDD-induced accumulation of bile acids in the liver hypothesized to promote biliary hyperplasia (21)), may be evident in urinary metabolite profiles. This study showed a dose-dependent increase in urinary TMAO occurred in the absence of liver pathology due to the AHR-mediated FMO3 induction in hepatocytes and may promote steatosis and cardiovascular disease. 1MN was also altered by low/moderate TCDD doses below the threshold for moderate to severe steatohepatitis. Like TMAO and 1MN, urinary metabolites altered by ≥ 10 μg/kg TCDD were likely mediated following AHR activation in hepatocytes although the involvement of extrahepatic metabolism cannot be excluded in the absence of hepatocyte-specific AHR knockouts. Overall, this study adds to the growing list of hepatic pathways disrupted following AHR activation by TCDD and demonstrates TCDD dose-dependent changes in the mouse urinary metabolome.

## EXPERIMENTAL PROCEDURES

### Study Design

Post-natal day (PND) 25 C57BL/6NCrl males from Charles River Breeding Laboratories (Kingston, NY) were acclimatized for five days prior to treatment. Mice were housed in Innovive Innocages (San Diego, CA) containing ALPHA-dri bedding (Shepherd Specialty Papers, Chicago) at an ambient temperature of 21° C with 30-40 % humidity and a 12 h/12 light/dark cycle. All mice were fed the TEKLAD diet 8940 (Madison, WI) *ad libitum*. At PND 30, mice were orally gavaged with 0.1 ml sesame oil vehicle (Sigma-Aldrich, St. Louis, MO) or 0.03, 0.1, 0.3, 1, 3, 10, or 30 μg/kg TCDD (AccuStandard, New Haven, CT) every 4 days for 28 days from Zeitgeber (ZT) 0 – 3 for a total of 7 administered doses. On PND 55, urine and feces were collected from individual mice, snap-frozen in liquid nitrogen, and stored at -80° C. Three days later on PND 58, mice were euthanized by CO_2_. Blood, liver, kidney, and epididymal adipose tissues were collected. Whole livers were weighed, snap-frozen in liquid nitrogen, and stored at -80° C. Mice were monitored every day for fluctuations in body weight. Mice were monitored every day for changes in chow weight. Food consumption per day was calculated by dividing the difference in chow weight from the previous day by the number of mice per cage.

### Histopathology

A liver sample was fixed in 10% formaldehyde for 24 hours before being transferred to a 30% ethanol solution. Liver tissue was sectioned at 5 microns and stained with hematoxylin and eoson (H&E). Whole glass histology slides were digitally scanned using an Olympus VS200 Research Slide Scanner. Liver sections were microscopically examined by a board-certified veterinary pathologist for TCDD-associated histopathology and semi-qualitatively scored for severity of inflammation, apoptosis or necrosis, lipid vacuolation, bile duct hyperplasia, and hepatocellular hypertrophy (0 = 0 % of liver tissue area ─ minimal (1) = 1 < 10 % of liver tissue area; mild (2) = 10 – 25 % of liver tissue area; moderate (3) = 26 – 50 % of liver tissue area; marked (4) = 51 – 75 % of liver tissue area; severe (5)= 76 – 100 % of liver tissue area affected). The median severity score for each dose was calculated. Severity scores were statistically tested by a Kruskal-Wallis followed by a Dunnett test in R (v4.2.1).

### 1H NMR Preparation, Acquisition, Analysis

Urine samples (500 μl) in a 1.5 mL Eppendorf tube were centrifuged at 12,000 *g* for 10 minutes at 4° C to remove debris. The supernatant (300 μL) was transferred to a clean 1.5 mL Eppendorf tube containing 35 μL of D_2_O and 15 μL of buffer (11.667 mM DSS [disodium-2,2-dimethyl-2-silapentane-5-sulphonate], 730 mM imidazole, and 0.47 % NaN_3_ in H_2_O). Samples (350 μL) were then transferred to a standard 3 mm thin-walled glass NMR tube for ^1^H NMR spectral analysis. All ^1^H NMR spectra were randomly collected on a Bruker Ascend HD 600 MHz spectrometer equipped with a 5 mm TCI cryoprobe and acquired at 25° C using the modified version of the first transient of the Bruker noesy-presaturation pulse sequence providing a high degree of quantitative accuracy. Spectra were collected with 128 transients and 16 steady-state scans using a 5-second acquisition time and a 5.1-second recycle delay. Prior to spectral analysis, all FIDs were zero-filled to 128K data points, and line broadened by 0.5 Hz. The methyl singlet produced by a known quantity of DSS (1000 μM) was used as an internal standard for chemical shift referencing (set to 0 ppm) and for quantification. All ^1^H NMR spectra were processed and analyzed using Chenomx NMR Suite Professional software package version 8.3 (Chenomx Inc., Edmonton, CA). Prior to statistical analysis, all NMR spectra were manually inspected for technical faults. Analysis identified 107 unique metabolites in the mouse urine samples. The concentration of all metabolites were normalized to creatinine signal followed by probabilistic quotient normalization (PQN). Each metabolite distribution was tested for differences across the dose response by a Kruskal-Wallis test. P-values were adjusted by the Benjamini-Hochberg method. Sixteen metabolites were determined to be significant (adjusted p-value ≤ 0.05). *Post hoc*, each metabolite was subject to a Dunnett test. The 1-D ^1^H NMR based metabolomics data was deposited to the MetaboLights which is the first general-purpose, open-access repository for metabolomics studies (75) under the submission of MTBLS960.

### RNA extraction

RNA was extracted as previously described (13, 31). Briefly, ∼100 mg of frozen tissues was submerged in 1.3 mL TRIzol and disrupted with a Mixer Mill 300 and a 3 mm stainless steel ball. Following the addition of chloroform, samples were manually shaken and left at room temperature for 3 minutes before centrifugation (12000 x *g*) at 4° C for 15 minutes. The supernatant was removed to another Eppendorf tube. Then, 125:24:1 phenol:chloroform:isoamylalcohol was added, and the mixture was centrifuged at 4° C for 10 minutes. The supernatant was removed, and equal parts isopropanol was added and left at -20° C overnight. The next day, the samples were centrifuged at 4° C for 15 minutes (12000 x *g*) and the isopropanol was removed. The pellet was washed with 70 % ethanol and centrifuged again for 10 minutes at 4° C (12000 x *g*). The ethanol was removed, and the pellet was broken up, dissolved in RNA Storage Solution (Thermo Scientific, Waltham, MA), heated at 70° C for 1 minute, and stored at -80° C. Total RNA was assayed for concentration and purity by a Nanodrop and examined for integrity using an Agilent 2100 Bioanalyzer.

### Bulk RNA sequencing, processing, and analysis

Bulk liver (n = 5 per dose) and kidney (n = 4-5 per dose) RNA-seq were performed by Novogene on a NovaSeq 6000 for a total of 20M 150 base-pair un-stranded paired-end reads per sample. Alignment to GRCm39 (release 104) was performed with STAR (v2.7.3a) (76) after trimming adaptors with Trimmomatic (v0.39-Java-11) (77). Quality control of sequencing and alignment data was respectively performed with fastqc (v0.11.7-Java-1.8.0_162) and qualimap (78) using R (v4.0.3). Separately, gene counts from liver and kidney were variance stabilized using DESeq2 (79) and modeled using Wolfinger’s mixed linear approach (80). Then, posterior probabilities (P1(t)’s) were calculated from the derived t-values using a Bayesian approach to reduce false discoveries. Genes were considered differentially expressed when the absolute fold-change was ≥ 1.5 and the P1(t) was ≥ 0.8. Data wrangling and transformation were done with R (v4.2.1). Principal component analyses were performed using the method ’prcomp’. Plots were generated through ggplot2 and formatted using Adobe Illustrator (v25.2.3). Raw and processed data for the liver and kidney are available at GSE203302 and GSE272652, respectively (28).

### Benchmark Dose (BMD) Analysis

BMD response values were calculated for bulk liver gene expression and urinary metabolites using BMDExpress2 (v2.3) (49). Each metabolite was fit to parameterized models: Exponential 2, Exponential 3, Linear, Polynomial 2, Polynomial 3, Hill, and Power. Fitting converged over a maximum of 250 iterations to calculate a 10 % BMD response value with a 95 % confidence level, consisting of a BMD upper (BMDU) and lower (BMDL) bound. Constant variance was assumed. Power was not restricted. BMDL and BMDU were utilized in the best model selection. A nested likelihood ratio test determined the best linear or polynomial model with a p-value cutoff of 0.05. Subsequently, the Akaike information criterion (AIC) was compared among the best linear or polynomial model and the remaining models. Hill models were flagged with less than 1/3^rd^ of the lowest positive dose. If a Hill model was flagged, the next best model according to AIC was selected if the p-value was greater than 0.05.

### Putative dioxin response element (pDRE) Identification

pDREs were identified as previously described (24, 31, 81). Briefly, the mouse genome (mm10 GRCm38 build) was computationally searched for the DRE core consensus core 5’-GCGTG-3’. Each core was extended by 7 base pairs (bp) upstream and downstream. The resulting 19 bp sequences were scored using a position weight matrix constructed from *bona fide* functional DREs. For annotation at the gene level, pDRE locations were compared against the regulatory region (10 kb upstream of the TSS together with 5’ and 3’ UTRs) and the coding sequence of each mouse gene obtained from the UCSC genome browser. The raw bedGraph file for the mouse pDRE analysis is available on Harvard Dataverse (82).

### Hepatic AHR ChIP-sequencing

ChIP-seq was performed as previously described (24) and is available on GEO (GSE97634). Briefly, liver samples were collected from C57BL/6NCrl adult male mice 2 hours after a single oral gavage with 30 μg/kg TCDD. Cross-linked DNA was immunoprecipitated with either rabbit IgG or rabbit IgG and anti-AHR as previously described (81, 83). Libraries prepared using the MicroPlex kit (Diagenode) were pooled and sequenced at a depth of approximately 30 M on an Illumina HiSeq 2500 at the Michigan State University (MSU) Research and Technology Support Facility Core. Read processing and analysis were performed using the MSU High-Performance Computing Center. Quality was determined using FASTQC v0.11.2 and adaptor sequences were removed using Cutadapt v1.4.1 while low-complexity reads were cleaned using FASTX v0.0.14. Reads were mapped to GRCm38 (release 76) using Bowtie v2.0.0 and alignments were converted to SAM format using SAMTools v0.1.19. Normalization and peak calling were performed using CisGenome (84) determined by comparison of IgG control and AHR enriched samples (n = 5) using a bin size (b) of 25 and boundary refinement resolution (bw) of 1 with default parameters.

### Metagenomics

Metagenomic analysis was performed as previously described (85). Briefly, microbial DNA from cecum contents (∼25 mg) of C57BL/6NCrl male mice repeatedly gavaged every 4 days for 28 days was extracted using the FastDNA spin kit for soil and sequenced on a NovaSeq 6000 by Novogene. Reads aligning to the C57BL/6NCrl genome were removed. The HUMAnN 3.0 bioinformatic pipeline was used with default settings to classify reads to UniRef90 protein identifications, which were mapped to enzyme commission (EC) number entries. Read abundance was normalized to gene copies per million reads (CPM). Multivariate association between dose and EC number relative abundance was established using Maaslin2 with the following settings: normalization (total sum scaling), analysis method (general linear model), and Benjamini-Hochberg multiple test correction. Raw sequencing files and processed data can be found at the NCBI Sequence Read Archive under the accession ID PRJNA719224.

### Western Blotting

Lysates (20 μg) and the PageRuler Prestained Protein Ladder (Thermo Scientific, Waltham, MA) were resolved via 10 % SDS-PAGE gels (Bio-Rad, San Diego, CA) and transferred to nitrocellulose membranes (GE Healthcare, Chicago, IL) using the Mini Trans-Blot Cell Unit (Bio-Rad, San Diego, CA) by wet electroblotting (100 V, 45 min). The membranes were then blocked with 5 % nonfat milk (in Tris-buffered saline [TBS]+0.01% Tween) for 1 hour and incubated with primary antibodies: anti-FMO3 (1:5000; ab126711; Abcam, Waltham, MA), anti-GLO1 (1:1000; MA531148, Life Technologies Corporation, Carlsbad, CA), anti-HAO1 (1:5000; ab194790; Abcam, Waltham, MA), anti-AGXT (1:1000; ab178708; Abcam, Waltham, MA), and anti-ACTB (1:1000; #4970; Cell Signaling, Danvers, MA) overnight at 4° C. Blots were visualized using horseradish peroxidase (HRP)-linked secondary antibodies of goat anti-mouse (1:10000; Elabscience, Houston, TX) and anti-rabbit (1:1000, Cell Signaling, Danvers, MA) and an ECL kit (Millipore Corporation, Billerica, MA). Membranes were scanned on a Sapphire Biomolecular Imager (Azure Biosystem, Dublin, CA). Protein density values were assessed and calculated using ImageJ (v1.53). Protein expression was standardized to ACTB levels per sample. Raw images are available at the Data Dryad data repository, including raw images with protein size ladders (86).

### Bulk TCDD exposure RNA-seq

Time-course liver (GSE109863), dose-response duodenum (GSE87542), dose-response jejunum (GSE90097), dose-response proximal (GSE171942) and distal (GSE89430) ileum, dose-response colon (GSE171941), dose-response femur (GSE104551), and dose-response gonadal white adipose tissue (GSE272683) RNA-seq data were processed as previously described (21, 87). Liver (n = 5) data were from C57BL/6NCrl males orally gavaged with sesame oil vehicle or 30 μg/kg TCDD and euthanized 2, 4, 8, 12, 24, 72, or 168 hours post-gavage. Duodenum (n = 3), jejunum (n = 3), proximal (n = 3) and distal (n = 3) ileum, femur (n = 3), colon (n = 3), and kidney (n = 4-5) data were from C57BL/6NCrl males orally gavaged every 4 days for 28 days with TCDD or sesame oil vehicle. Gonadal white adipose tissue (GWAT) data were from C57BL/6NCrl females gavaged every 4 days for 92 days with TCDD or sesame oil vehicle. Unlike liver samples, intestinal and GWAT tissue collected from C57BL/6NCrl females were not collected within a narrow ZT (0–3) window.

### Single nuclear RNA-sequencing (snRNA-seq) analysis

Available at a public repository (GSE184506), TCDD dose-response snRNA-seq of liver cell types was processed as previously described (88). Male C57BL/6NCrl mice were gavaged with TCDD (0.03 – 30 μg/kg) or sesame oil vehicle every 4 days for 28 days. 11 Cell types were identified in line with previous studies (89–91) by cluster analysis and semi-automated annotation. Dot plots of average cell type-specific gene expression at a certain dose of TCDD or sesame oil vehicle were generated in R using Seurat.

### Publicly Available Bulk Liver Gene Expression Data Analysis of Certain Genetic Knockouts

Verification of TCDD-induced FMO3-induction-related gene expression was performed by pulling down bulk liver RNA-seq data from wild-type (WT) and *Fmo3* knockout C57BL/6 mice, exposed to TCDD (GSE191138). Mice were treated with weekly injections of 25 μg/kg TCDD for 6 weeks (41). The data was aligned and transformed identically to the TCDD dose-response bulk liver RNA-seq (See **Bulk RNA sequencing, processing, and analysis**). Verification of TCDD-induced, AHR-mediated gene regulation was performed by pulling down bulk liver microarray data from WT and *Ahr* knockout C57BL/6 mice, orally gavaged with a bolus of TCDD or corn oil vehicle (GSE15858). Data was processed in R as previously described (43).

### GC-MS

Extraction, derivatization, and measurement of hepatic metabolites by gas-chromatography mass spectrometry (GC-MS) were performed using protocols adapted from the publicly available Michigan State University Mass Spectrometry and Metabolomics Core website (https://rtsf.natsci.msu.edu/sites/_rtsf/assets/File/MSU_MSMC_004_v1_2_Two_phase_extraction_of_ metabolites_from_animal_tissues.pdf). Briefly, ∼50 mg of frozen liver tissue (*n* = 5 per dose) was supplemented with 6 nanomoles of ^13^C and ^15^N labeled amino acid internal standards (767964, Sigma-Aldrich) and lysed with a Mixer Mill 300 and a 3 mm stainless steel ball in a solution of methanol/chloroform (1:2 v/v), 1 % formic acid, and 0.01 % BHT. Subsequently, the samples were sonicated for 15 minutes. The mixture was twice supplemented with Milli-Q water, vortexed, centrifuged, and the resulting aqueous supernatant was transferred to a separate Eppendorf tube and made basic by the addition of NaOH. The samples were evaporated to dryness by a SpeedVac without heat and stored at -20° C. For derivatization, the dried samples were set at 60° C for 12 hours with 100 μL of 40 mg/mL of methoxyamine HCl dissolved in pyridine and then incubated again at 60° C for another 12 hours with an additional 100 μL of N-Methyl-N-tert-butyldimethylsilyltrifluoroacetamide (MTBSTFA) containing 1% tert-butyldimethylsilyl chloride. Derivatized samples (1 μL) were injected with a 1:10 split at 230° C and analyzed on an Agilent 7890A GC/ single quadrupole mass spectrometer with 5975C inert XL MSD. The carrier gas was helium, which flowed at 1.0 mL/min. Separation was achieved on an Agilent J&W VF-5ms (30 m x 0.25 mm x 0.25 μm) (Agilent, Santa Clara, CA) using the following temperature profile: initially set at 40° C, ramped to 80° C at 20° C/min, immediately ramped to 250° C at 5° C/min and held for 1 minute, and ramped to 320° C at 50° C/min and held for 5 minute. The mass spectrometer was operated at an electron ionization of 70 eV and a scan range of 50-600 *m/z*. Glycolic acid (PHR1427, Sigma), hydroxyproline (PHR1939, Sigma), oxalic acid (PHR3291, Sigma), and urea (PHR1406, Sigma) were used as external standards to calculate metabolite levels per sample. Quantification of the derivatized metabolites and internal standard were calculated using the peak area from the extracted ion chromatogram. The SIM ion used for derivatized glycolic acid, hydroxyproline, oxalic acid, and urea were 247, 416, and 261m/z, respectively. Derivatized glycolic acid and hydroxyproline were normalized to labeled glycine, while oxalic acid was normalized to labeled alanine. Oxalic acid and alanine were quantified by separate runs in the SIM mode, while the other compounds were quantified from runs in the scan mode. Additionally, hydroxyproline was normalized to labeled proline. After calculating concentrations via a linear calibration curve, concentrations were normalized to the liver weight per sample from which metabolites were extracted. The raw and processed files were uploaded to metabolomics workbench (ST003476).

### Activity Assay

GLO1 activity was assayed using a commercial colorimetric kit (ab241019; Abcam, Waltham, MA) according to the manufacturer’s instructions. Liver tissue was disrupted with a Mixer Mill 300 and a 3 mm stainless steel ball.

## Supporting information

Supplementary Tables 1 through 3

## DATA AVAILABILITY

All sequencing, metabolomic, and Western blot data have been deposited in publicly available repositories. Accession numbers and the names of the data repositories can be found in the data-specific experimental procedures sections.

## SUPPORTING INFORMATION

This article contains supporting information.

## ACKNOWLEDGEMENTS

GC-MS method development and instrumentation was performed in collaboration with the MSU mass spectrometry and metabolomics core, especially Dr. Cassandra Johnny.

## AUTHOR CONTRIBUTIONS

W. J. S., S. F. G., T. Z. conceptualization; W. J. S., R. R. F., W. J. S., A. Y., D. G., and R. N. investigation; W. J. S. visualization; W. J. S., writing-original draft; J. R. H. formal analysis; W. J. S., R. R. F., A. Y., R. N., J. R. H., S. F. G. and T. Z. writing-review and editing; W. J. S. project administration; W. J. S. and A. Y. methodology; T. Z. and S.F.G. funding acquisition.

## FUNDING AND ADDITIONAL INFORMATION

This project was supported by the National Institute of Environmental Health Sciences (NIEHS) Superfund Research Program [NIEHS SRP P42ES004911] and the NIEHS Research Project Grant Program [NIEHS R01ES029541] to T. Z. W. J. S. and R. R. F. were supported by NIEHS Multidisciplinary Training in Environmental Toxicology [NIEHS T32ES007255]. The content is solely the responsibility of the authors and does not necessarily represent the official views of the National Institutes of Health.

## ABBREVIATIONS

DLC: dioxin-like compound
TCDD: 2,3,7,8-tetrachlorodibenzo-*p*-dioxin
AHR: aryl hydrocarbon receptor
TMA: trimethylamine
TMAO: trimethylamine *N*-oxide
1MN: 1-methylnicotinamide
snRNA-seq: single-nuclear RNA-seq
MASLD: metabolic dysfunction associated steatotic liver disease
PCDD: polychlorinated dibenzo-p-dioxins
HCC: hepatocellular carcinoma
SLD: steatotic liver disease
BHLH: basic-helix-loop-helix
PAS: Per-Arnt-Sim
ARNT: AHR nuclear translocator
PND: post-natal day
ZT: Zeitgeber
DRE: dioxin response element
H&E: hematoxylin and eosin
BMD: benchmark dose
BMDU_10%_: BMD upper bound with a 10% confidence interval
BMDL_10%_: BMD lower bound with a 10% confidence interval
PQN: probabilistic quotient normalization
CN: creatinine normalization
PCA: principal component analysis
PEV: percent explained variance
AGE: advanced glycation end-products
GC-MS: gas chromatography mass spectrometry
PtdC: phosphatidylcholine
DMA: dimethylamine
TSS: transcription start site
BCAA: branched-chain amino acids
BCKA: branched-chain α-keto acids
BCKDH: branched-chain alpha-keto acid dehydrogenase
NAD+: nicotinamide adenine dinucleotide
ALT: alanine aminotransferase
CPM: counts per million
FMO3: flavin-containing monooxygenase family member 3
EC: enzyme commission

---

**Supplementary Figure 1.**
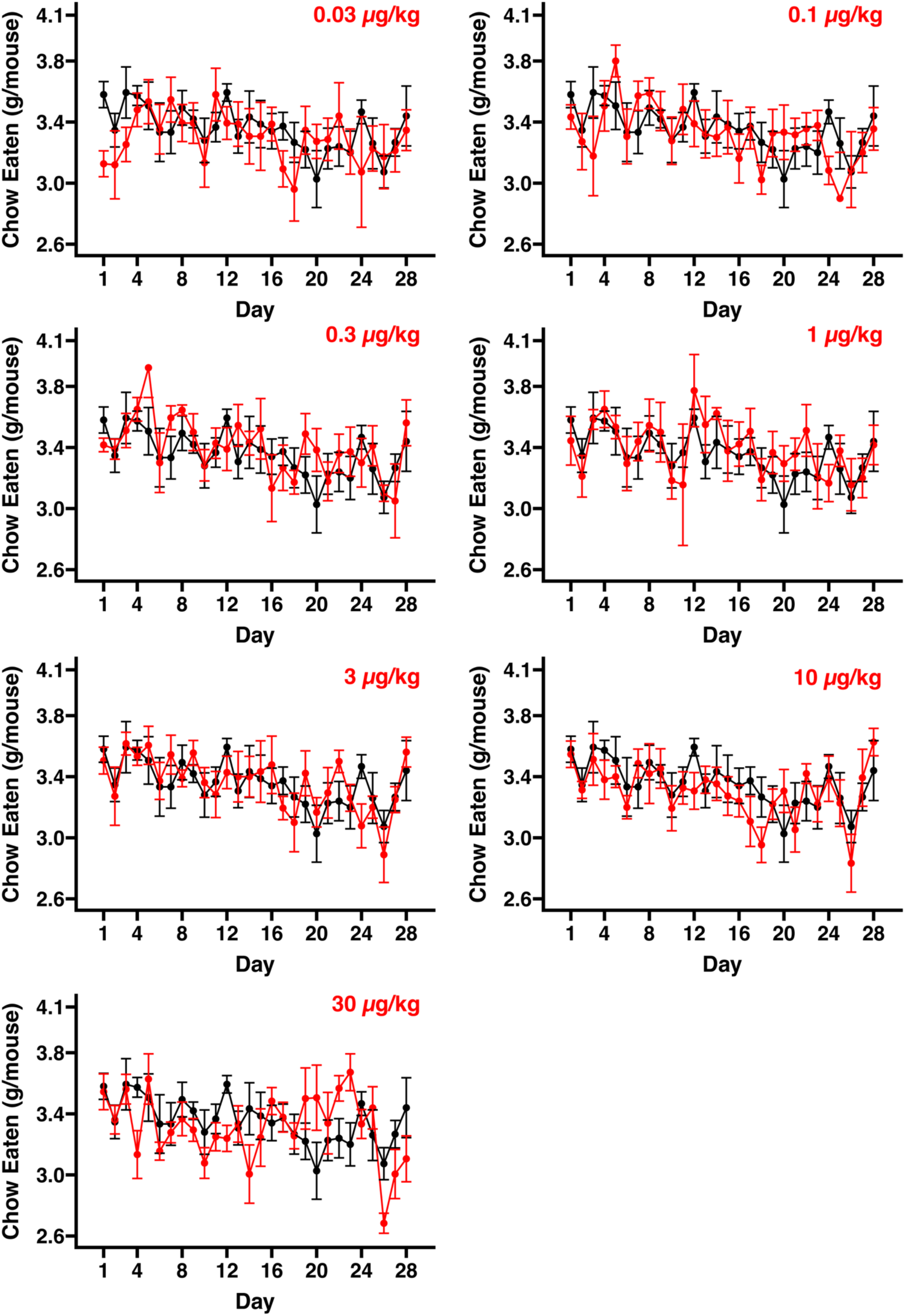
Chow consumption of mice gavaged with TCDD every 4 days for 28 days. Consumed chow per cage (n = 5-6 cages per dose) was monitored by weighing food each day of the study between ZT0-3. The black line in each plot indicates the average chow eaten per mouse per day following sesame oil (vehicle control) gavage. The red lines indicate the average chow eaten per mouse per day following gavage with TCDD. The error bars indicate standard error. A repeated measures ANOVA test was used to detect chow consumption differences with respect to dose. Student’s t-test was used to assess pairwise differences in food consumption per mouse per day between sesame oil and TCDD gavaged mice. P-values from all days for each pairwise comparison were adjusted using the Benjamini-Hochberg method. No significant (p-value ≤ 0.05) difference in chow consumption was detected at any dose.

**Supplemental Figure 2.**
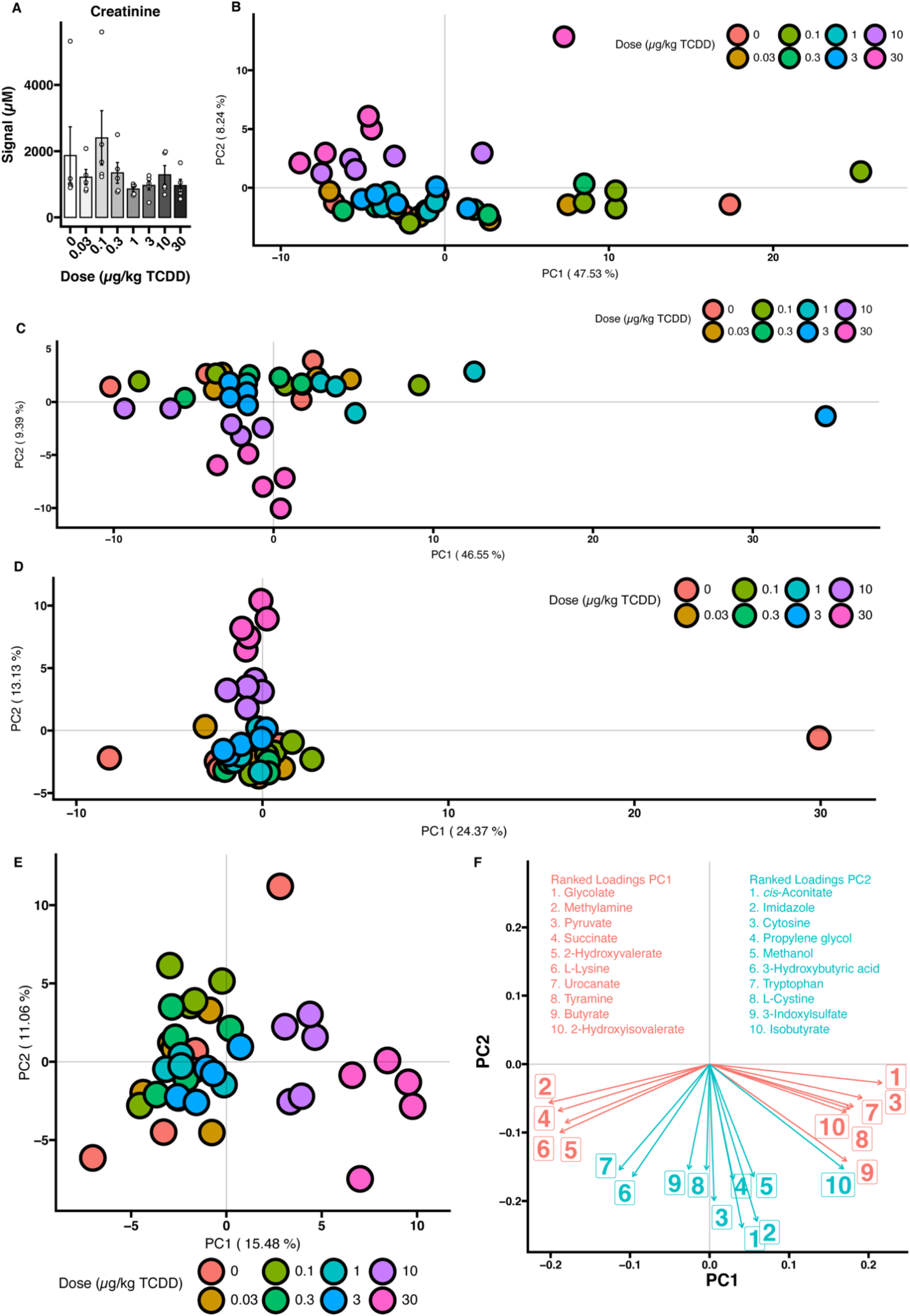
1-D ^1^HNMR normalization. (A) Un-normalized creatinine signal. Difference between dosing groups was tested by a Kruskal-Wallis test. Post hoc, groups were tested by Dunnett’s test for difference with sesame oil treated mice. No significance (p-value ≤ 0.05) was detected. Principal components 1 and 2 from PCA of 1-D ^1^HNMR after (B) no normalization, (C) creatinine normalization (CN), (D) probabilistic quotient normalization (PQN), and (E) CN followed by PQN. (F) The most important metabolites (*i.e.*, loadings) for driving the separation of samples by PCs 1 and 2 from (E).

**Supplementary Figure 3.**
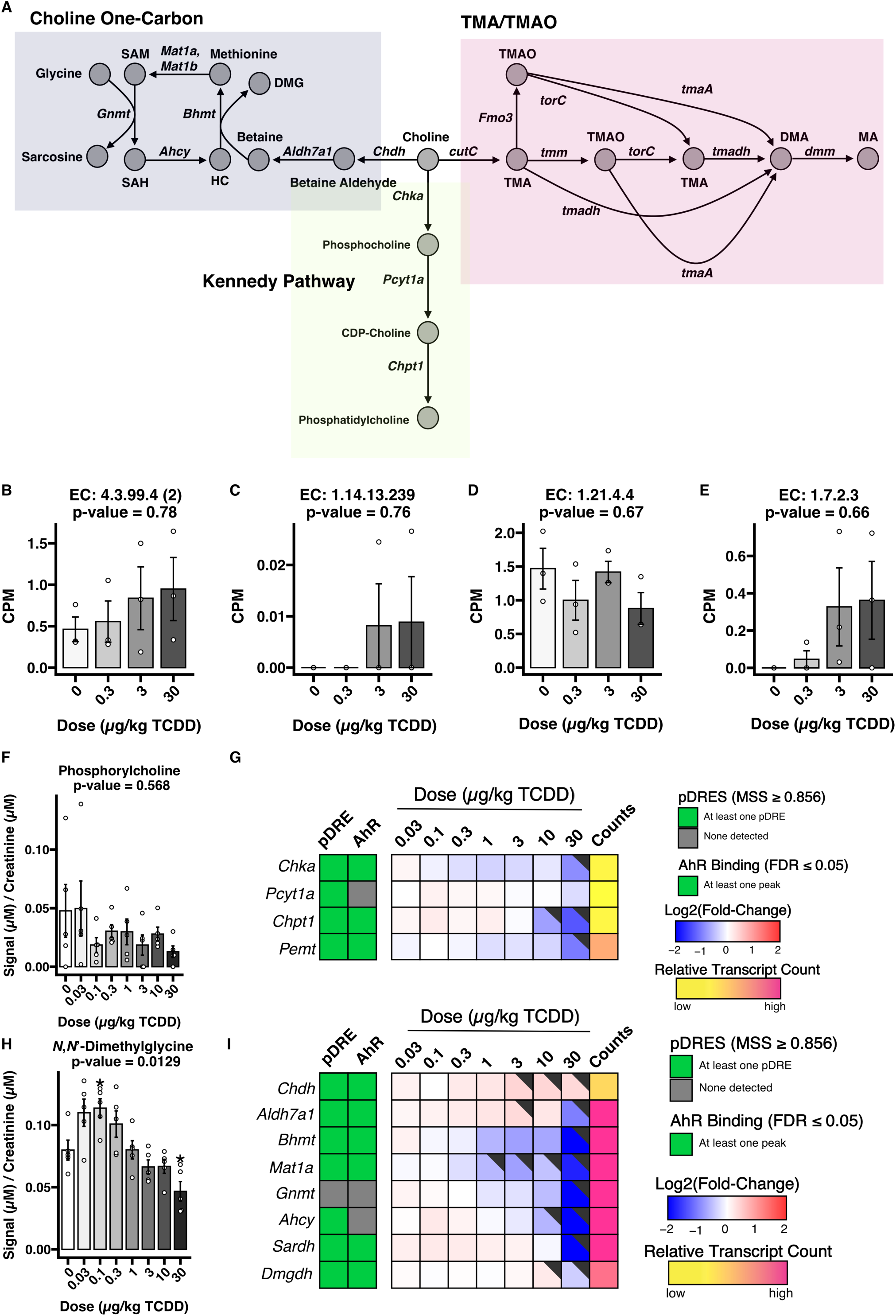
Possible metabolic routes of dietary choline. (A) Choline metabolism pathway. Circles indicate metabolites, while arrows indicate an enzymatic reaction encoded by a particular gene. (B – E) Metagenomic copies per million (CPM) for all detected TMA/TMAO pathway genes. (F) Urinary phosphorylcholine levels as measured by 1-D ^1^HNMR (n = 5). An asterisk (*) indicates significance from a *post hoc* Dunnett test (p ≤ 0.05). (G & I) Heatmaps of phosphatidylcholine and betaine biosynthesis. pDREs were determined by a position weight matrix using a Matrix Similarity Score (MSS) cut off of ≥0.856 based on the sequence of characterized functional DREs. Hepatic AHR ChIP-seq was detected in mouse two hours after oral gavage with 30 μg/kg TCDD (GSE97634). The green tiles indicate an FDR of ≤ 0.05. In the dose response bulk RNA-seq gene expression, the black flags in the upper right hand corner of a tile indicate a P1(t) of at least 0.8. The Counts column represents the maximum estimate of aligned reads of each gene. Lower levels of expression (≤500 reads) are depicted in yellow with higher level sof expression (≥10,000) depicted in pink. (H) Urinary *N,N’*-dimethylglycine levels were measured by 1-D ^1^HNMR (n = 5). An asterisk (*) indicates significance from a *post hoc* Dunnett test (p ≤ 0.05). Abbreviations: SAM = S-adenosyl methionine; SAH = S-adenosyl homocysteine; HC = homocysteine; DMG = N,N’-dimethylglycine; CDP-choline = cytidine diphosphate-choline; TMA = trimethylamine; TMAO = trimethylamine N-oxide; DMA = dimethylamine; MA = methylamine.

**Supplementary Figure 4.**
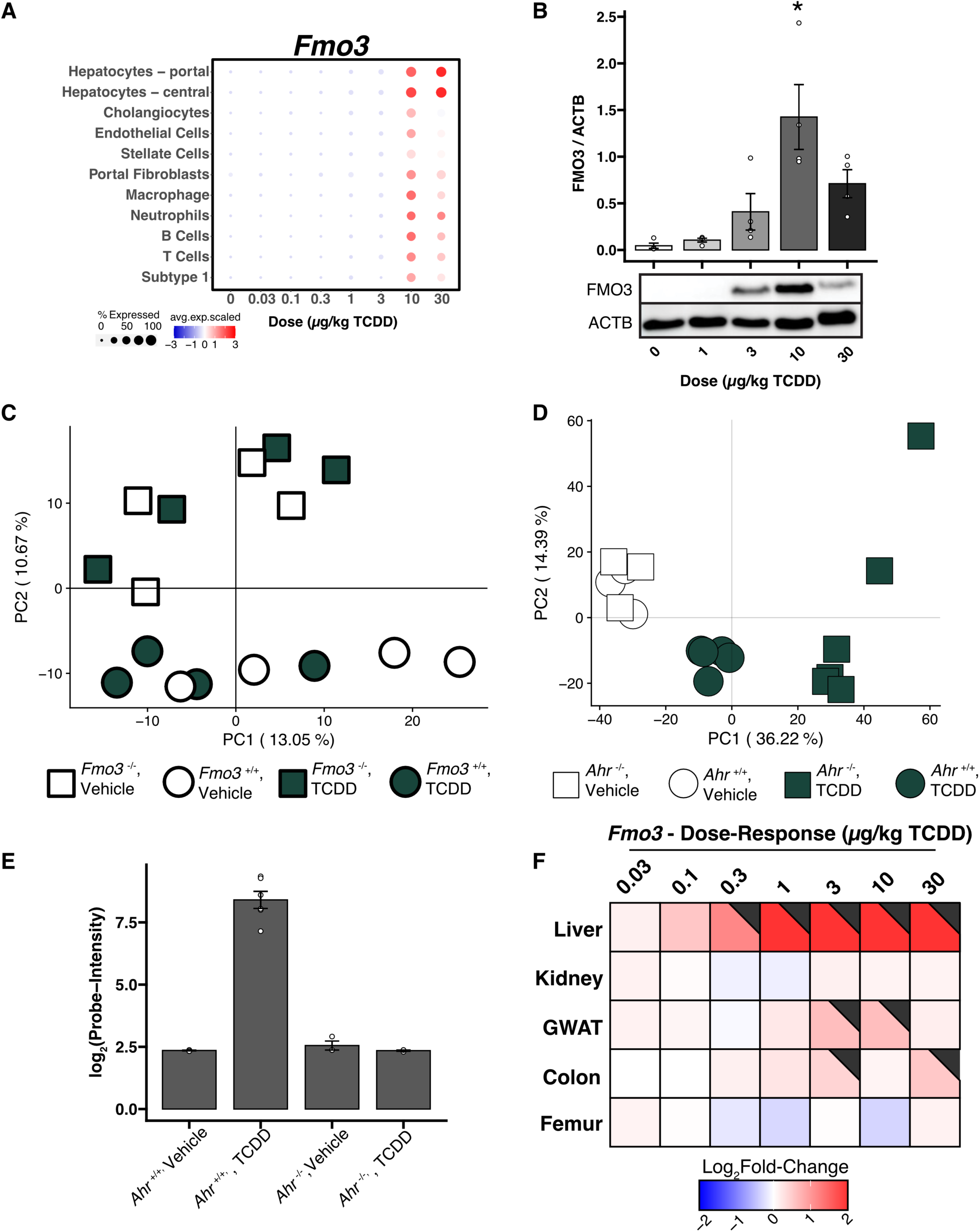
Murine genetic knockout models and TCDD-induced TMAO production. (A) snRNA-seq analysis of *Fmo3* expression in individual hepatic cell types from mice gavaged every 4 days for 28 days with TCDD (n = 3). The size of the dot represents the percent of cells that expressed *Fmo3*. The color of the dot is the average expression level of *Fmo3*, centered and scaled across all cells. (B) Western blot of FMO3 in liver cell lysates (n = 4). Densitometry was performed using ImageJ. A Kruskal-Wallis test was performed, followed *post hoc* by a Dunnett’s test with respect to sesame oil vehicle control. An asterisk (*) indicates significance (p ≤ 0.05). (C) Principal components 1 and 2 of RNA-seq data (GSE191138) from wild type and AHR knockout mouse models treated with vehicle and TCDD RNA-seq (Massey *et al.*). Data was pulled down from the sequencing read archive (SRA) and aligned to GRCm39 (release 104) with STAR (v2.7.3a). Data was variance stabilized by DESeq2 before principal component analysis. (D) Principal components 1 and 2 of microarray data (GSE15858) from wild type and AHR knockout mouse models treated with vehicle and TCDD (Boutros *et al*.). Microarray data was pulled down from the gene expression omnibus, and normalization of signal intensity per probe was performed using the R package limma. (E) *Fmo3* gene expression from microarray data (GSE15858) from wild type and AHR knockout mouse models treated with vehicle and TCDD (Boutros *et al*.). Bars show means, and error bars indicate standard error. (F) *Fmo3* gene expression across multiple tissues from mice gavaged with TCDD over 28 or 92 days.

**Supplementary Figure 5.**
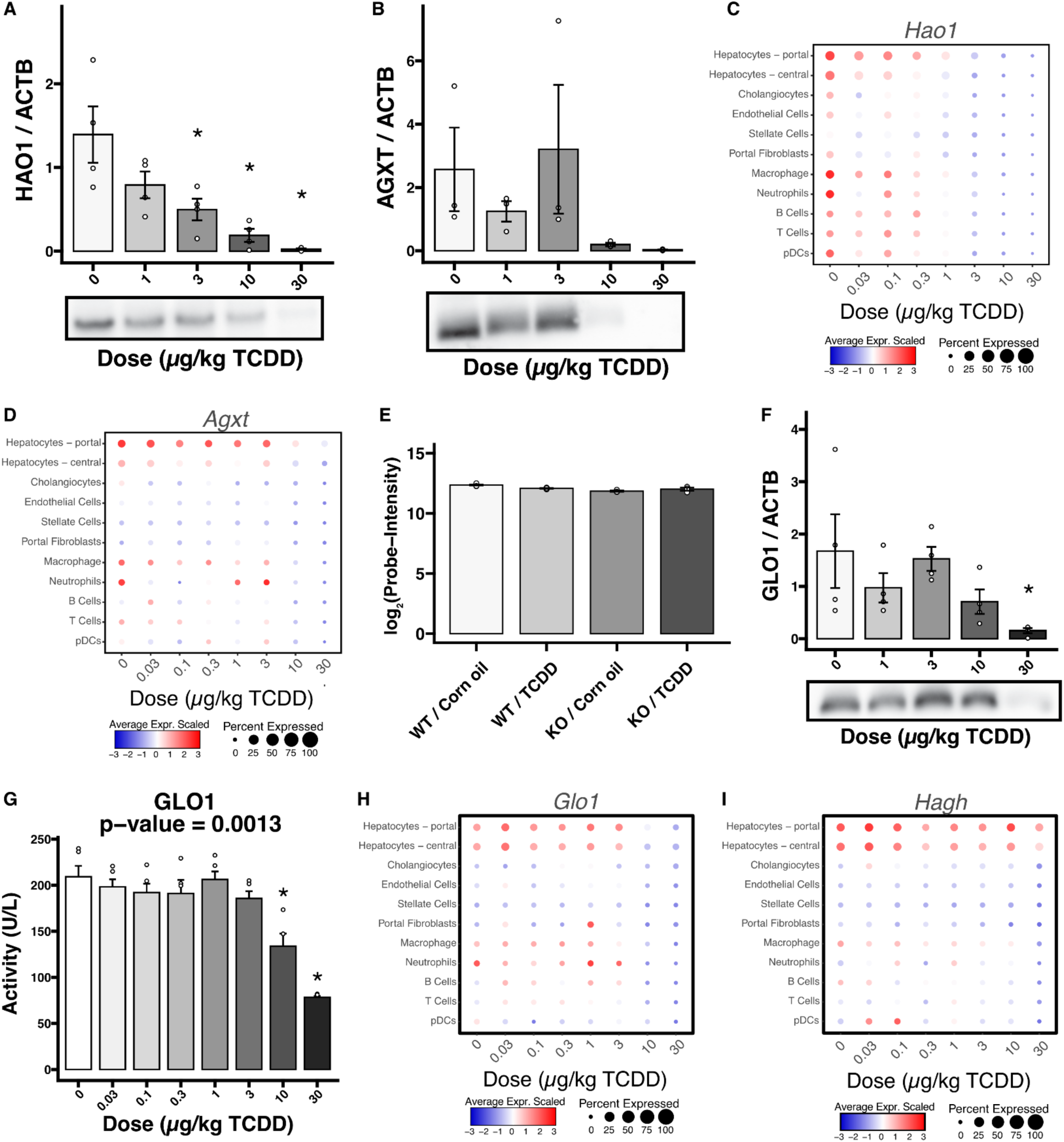
Hydroxyproline and glyoxal metabolism. (A – B) Densitometry analysis of Western blots of HAO1 and AGXT (n = 3 – 4). Bars indicate mean, and error bars indicate standard error. P-values were calculated using a Kruskal-Wallis test. An asterisk (*) indicates significance from a *post hoc* Dunnett test (p-value ≤ 0.05). (C – D) Average scaled expression and percent expressing cells of *Hao1* and *Agxt* from liver snRNA-seq (GSE184506). (E) *Hao1* gene expression of microarray data (GSE15858) from wild type and AHR knockout mouse models treated with vehicle and TCDD (Boutros *et al*.). Bars show means, and error bars indicate standard error. (F) Densitometry analysis of Western blots of GLO1 (n = 4). Bars indicate mean, and error bars indicate standard error. P-values were calculated using a Kruskal-Wallis test. An asterisk (*) indicates significance from a *post hoc* Dunnett test (p-value ≤ 0.05). (G) GLO1 activity as measured by a colorimetric kit (ab241019) in hepatic lysate (n = 5). Bars indicate mean, and error bars indicate standard error. P-values were calculated using a Kruskal-Wallis test. An asterisk (*) indicates significance from a *post hoc* Dunnett test (p-value ≤ 0.05). (H – I) Average scaled expression and percent expressing cells of *Glo1* and *Hagh* from liver snRNA-seq (GSE184506).

**Supplementary Figure 6.**
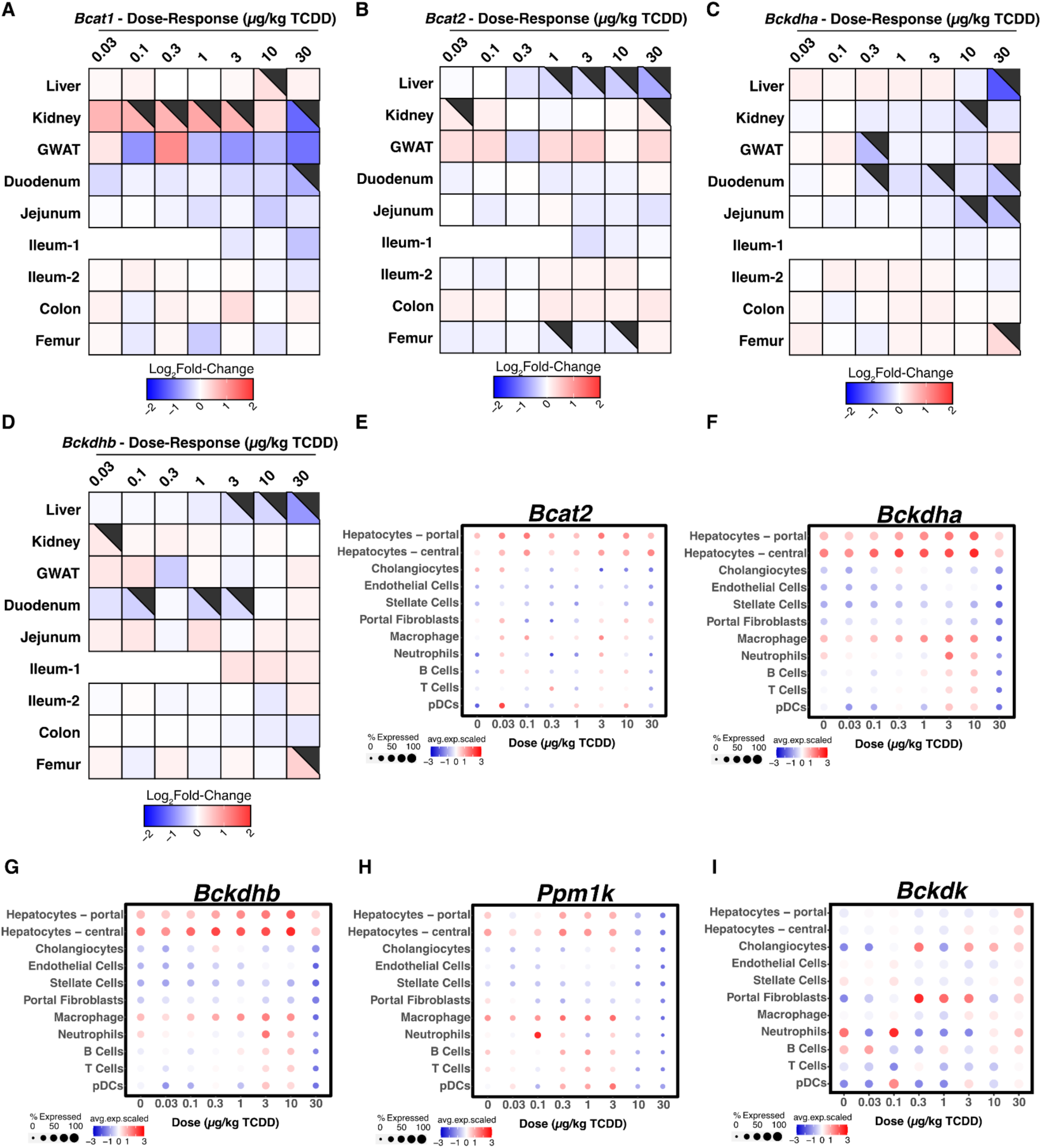
Alternative Branched-Chain Amino Acid Catabolism. (A - E) Average scaled expression and percent expressing cells of select genes from liver snRNA-seq (GSE184506). (F) Heatmap of hepatic BCAA catabolism related genes. pDREs were determined by a position weight matrix with a Matrix Similarity Score (MSS) ≥ 0.856. AHR ChIP-seq was detected in mouse liver two hours after oral gavage with 30 μg/kg TCDD – green tiles indicate an FDR of ≤ 0.05 (GSE97634). In the dose response bulk RNA-seq gene expression, the black flags indicate a P1(t) ≥ 0.8. The Counts column refers to the maximum estimate of aligned reads of each gene where a lower level of expression (≤ 500 reads) is depicted in yellow and a higher level of expression (≥ 10,000) is depicted in pink.

**Supplementary Figure 7.**
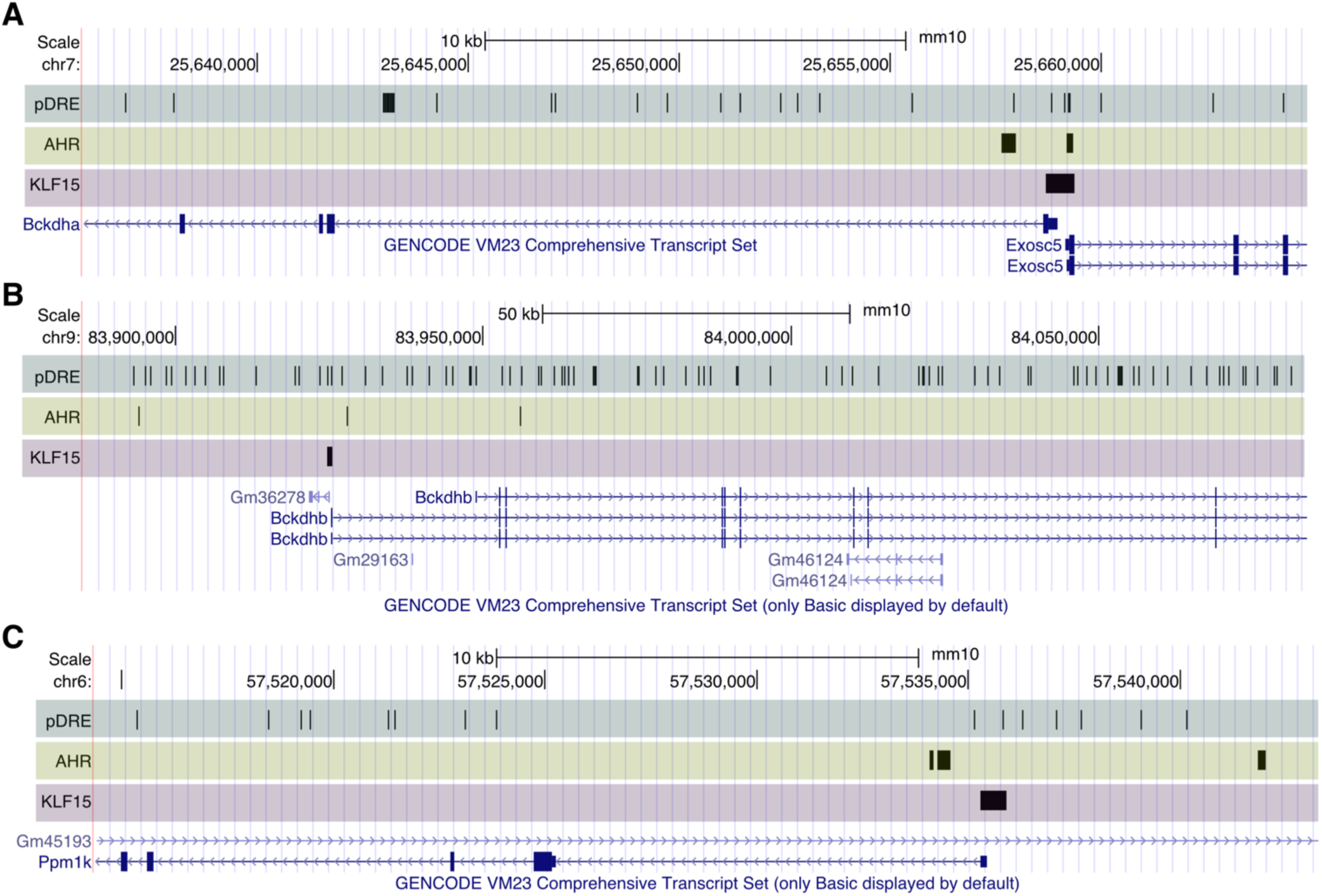
UCSC genome tracks of (A) *Bckdha*, (B) *Bckdhb*, and (C) *Ppm1k*, respectively. Arrows point toward 3’ end. The putative dioxin response elements (pDRE) were determined by a position weight matrix with a Matrix Similarity Score (MSS) ≥ 0.856. The aryl hydrocarbon receptor (AHR) track indicates binding two hours after mice were administered a single oral gavage of 30 μg/kg TCDD (GSE97634). The Krüppel-like factor 15 (KLF15) track indicates KLF15 binding in mice at homeostasis (GSE166083).

